# Adhesive silk hydrogel patches for localized and sustained delivery of cell-derived nanovesicles

**DOI:** 10.64898/2026.05.03.722555

**Authors:** Auriane Drack, Alin Rai, Hien A Tran, Jelena Rnjak-Kovacina, David W. Greening

## Abstract

The transplantation of stem cell-derived extracellular vesicles (EVs) holds promise for tissue repair and regeneration, but scalable production and effective delivery to target tissue remain major challenges. Here, we present a biomaterial platform that combines high-yield, scalable nanovesicles (NVs) – EV mimetics derived from human induced pluripotent stem cells – with an adhesive silk hydrogel patch for localized and sustained delivery. We show that this platform enables efficient NV encapsulation via visible light crosslinking and supports controlled release over short (2 days), intermediate (7 days), and extended (up to 28 days) periods, while maintaining adhesion to heart tissue. Importantly, the sustained delivery of NVs for 3 days *in vitro* results in promoting anti-fibrotic cell remodeling and significant functional recovery of primary myofibroblast activation, modulating integrin signaling, actomyosin organization, and cell-matrix adhesion networks. Finally, we demonstrate biocompatibility, retention, and anti-fibrotic function of the patch in a murine ischemia-reperfusion injury model. Thus, we establish the proof-of-principle that di-tyrosine silk hydrogels can be used as a strategy to encapsulate and deliver NVs to the heart, thus offering an innovative delivery platform for NVs.

**Statement of significance:** Extracellular vesicles (EVs) represent an emerging frontier in tissue engineering. Their cell-specific cargo contains biological information capable of repairing and regenerating injured tissues. However, their clinical translation is hindered by limited manufacturing scalability, undefined dosing and modes of administration, and low organ retention, particularly in the heart. This study addresses these challenges by combining stem cell–derived nanovesicles (NVs), which mimic biological EVs, with an adhesive hydrogel patch for localized and sustained delivery to the heart. We provide proof-of-principle that di-tyrosine photo-crosslinked silk hydrogels are a suitable delivery platform for cell-derived NVs, preserving NV bioactivity and their ability to remodel recipient cells following delivery both *in vitro* and *in vivo*. This study integrates three key advantages: (i) the use of scalable iPSC-derived nanovesicles as an EV-mimetic platform, addressing limitations in EV manufacturing; (ii) a mechanically robust and tunable silk fibroin hydrogel formed via visible light-induced di-tyrosine crosslinking without chemical modification; and (iii) an injection-free, adhesive patch-based delivery strategy enabling localized and sustained therapeutic administration to the heart. This innovative platform represents a significant advancement in the fields of nanomedicine and biomedical engineering.

Graphical abstract

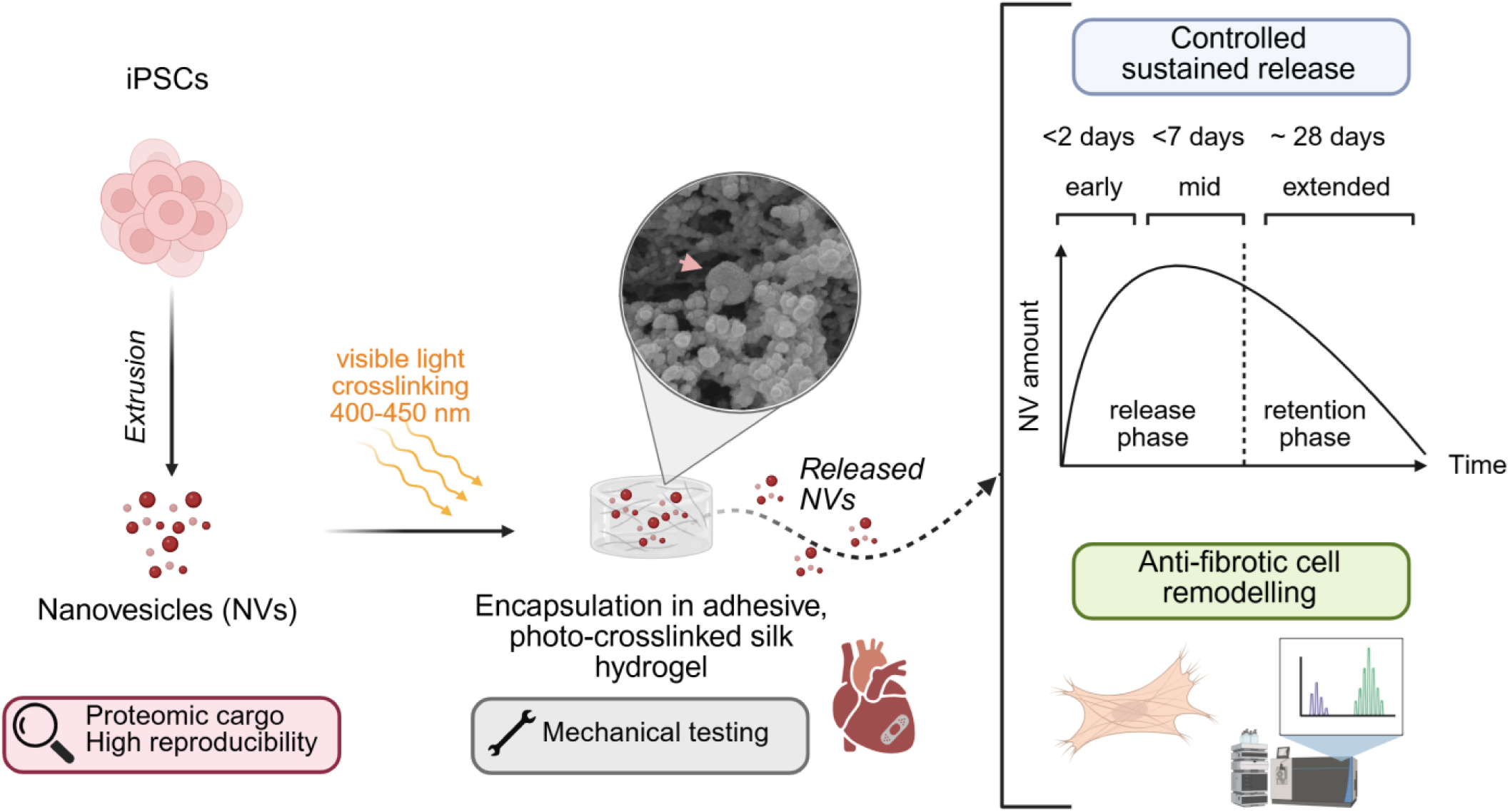

## 1. INTRODUCTION

Cell–cell communication is fundamental to the coordination of biological processes, playing a central role in development, homeostasis, and the response to physiological and pathological stimuli [1]. Among the various mechanisms by which cells exchange information, extracellular vesicles (EVs) have emerged as key mediators of intercellular communication [2]. EVs are lipid bilayer-enclosed particles naturally released by cells that transport a complex bioactive cargo to elicit signaling [3, 4], phenotypic changes, and functional responses in recipient cells [5–7]. EVs have emerged as promising cell-free therapies for their inherent properties such as immune modulation, biocompatibility, stability in circulation, tissue-targeting potential and their ability to act as natural carriers to protect, transport, and deliver bioactive factors across biological barriers [8]. EVs are currently being explored for a wide range of applications, including in cell and cancer therapy, wound healing, tissue repair and regeneration [9–17], including the heart [6, 18].

Despite the significant promise of EV-based therapies, major challenges hinder their clinical translation including their inherently low yield from stem cell source, lack of standardization in manufacturing process, challenges in understanding their composition as well as difficulties in demonstrating pharmacokinetics and efficacy [19]. To address the limitation of large-scale EV production, we and others have developed a pipeline to generate EV-like particles, termed cell-derived nanovesicles (NVs). NVs can be efficiently generated in large amounts by mechanically extruding cells rather than relying on the natural EV secretion [20]. Specifically, we demonstrated that NVs generated from stem cells are readily internalized by target cells, reduce fibroblast activation, enhance angiogenesis, and promote cardiomyocyte survival, thus underscoring their potential for tissue repair therapies [18, 21]. However, the retention time of both stem cell EVs and NVs in tissue by intravenous administration may be insufficient to achieve their full therapeutic potential [22], raising questions about how best to deliver cell-free therapies over prolonged periods.

Clinical application of acellular therapies depends on their efficient administration and delivery to target tissues, such as the heart [23, 24]. However, systemic administration of EVs often results in rapid clearance and poor cardiac retention [25], while direct intramyocardial injection is highly invasive and restricts dosing flexibility[17, 26]. To address these limitations, hydrogel-based delivery systems have emerged as transformative solutions for cardiac therapeutics, enabling localized delivery of drugs and therapeutic biomolecules including nucleic acids [27, 28], growth factors [29, 30], cell secretome [31], peptides [32, 33], and gene regulatory systems [34]. Notably, a recent meta-analysis of injectable hydrogel-based combination therapy for myocardial infarction reported that EV-loaded hydrogels produced the greatest functional improvement in cardiac outcomes compared with cells, drugs, or cytokines [35]. Other than direct intramyocardial injection [36], biomaterial-based therapies can also be delivered through pre-formed patches sutured onto the epicardium [37, 38], catheter-based delivery [39, 40], intrapericardial injection for *in situ* patch formation [41], and spray-painting techniques [42]. These strategies offer improved spatial control over therapeutic delivery, provide mechanical support to the injured myocardium, and may reduce procedural invasiveness. However, adhesive biomaterials capable of direct attachment to heart tissue and integration with the host myocardium for sustained therapeutic delivery remain largely unexplored [43, 44].

To overcome these challenges, silk fibroin has been utilized as a natural biomaterial for cardiac biomedical applications due to its biocompatibility, porosity, mechanical strength, and tissue-adhesiveness [45–48]. The therapeutic potential of silk fibroin has been demonstrated through diverse cardiac applications, including extracellular matrix-incorporated scaffolds that effectively mimic native cardiac tissue structure and function [49], injectable hydrogels that prevent adverse left ventricular remodeling post-myocardial infarction [50], and sustained delivery platforms for biomolecules, including EVs for vascular repair [51]. Developments in silk biofunctionalization, particularly di-tyrosine photo-crosslinking for covalent biomolecule immobilization, provide a biocompatible microenvironment for promoting cell function without chemical modifications to the base biomaterial [52].

This study presents the first report of combining iPSC-derived nanovesicles, which mimic EVs in their size, composition and function, with di-tyrosine crosslinked silk hydrogels for direct cellular and cardiac delivery. By combining detailed characterization with proteomic profiling of NVs and their remodelled target cells and organ following release from silk, we demonstrate this as a functional bio delivery platform for the treatment of cardiac fibrosis following MI.

## 2. MATERIAL AND METHODS

### 2.1 Cell culture

The human RM3.5 GT-Td Tomato iPSC line, kindly provided by Katerina Vlahos (Murdoch Children’s Research Institute, Melbourne), was maintained on growth factor reduced Matrigel-coated plates (Corning, #256231) in StemFlex medium (Gibco, A3349401) following the manufacturer’s protocol. Cells were passaged at confluency using an enzyme-free reagent (ReLeSR, Stem Cell Technologies). Media was replaced every two days. Fluorescent imaging to confirm Td-Tomato tag presence was performed using an inverted Olympus IX71 microscope at 20× magnification.

Human primary cardiac fibroblasts (ventricular, single donor; Sigma-Aldrich, CC-2904) were cultured on 1% gelatine-coated flasks (bovine skin Type B, Sigma-Aldrich, 9000-70-8). Cells were maintained at 37°C in 5% COL in a growth medium consisting of 75% Cardiac Fibroblast Growth Medium (Lonza, CC-4526), 25% DMEM/F12 (Gibco, 11320033), 10% 0.22 µm filtered fetal calf serum (Gibco, 10099141), and Penicillin-Streptomycin (Gibco, 15140122). Cells were passaged at a 1:3 ratio by surface area and used for experiments at passage 5–7.

Human umbilical vein endothelial cells (HUVECs, Angio-proteomie, cAP-0001) were cultured on 1% gelatin-coated flasks (bovine skin Type B, Sigma-Aldrich, 9000-70-8). Cells were maintained at 37°C in 5% COL in Endothelial Cell Growth Medium-2 BulletKit™ (Lonza, CC-3156 with CC-4176 supplements) prepared according to manufacturer’s instructions. Medium was changed every two days, and cells were passaged at confluency using 0.05% trypsin-EDTA (Gibco, 15400054) and reseeded at 1.0 × 10□ cells/cm² on 1% gelatin-coated plates. Cells were used for experiments at passage 5-7.

### 2.2 Silk hydrogel preparation

Bombyx mori silk cocoons were sourced from Sato Yama, Japan. Lithium bromide (LiBr), sodium carbonate (NaLCOL), sodium persulfate (SPS), Tris(2,2′-bipyridyl)dichlororuthenium(II) hexahydrate (Ru), and dialysis tubing (Snakeskin, 3500 MWCO) were obtained from Sigma-Aldrich. Phosphate-buffered saline (PBS) tablets and sodium chloride (NaCl) were purchased from Thermo Fisher Scientific.

To obtain regenerated silk fibroin, silk cocoons were processed as described [53]. In brief, 5 g of silk, cut into small fragments, was boiled in 2 L of 0.02 M Na□CO□ for 30 mins to remove sericin. The degummed silk fibers (0.25 g/mL) were dried and subsequently dissolved in 9.3 M LiBr at 60 °C for 5 h. To remove LiBr, the solution was dialyzed against MilliQ water for 72 h using Snakeskin tubing (3500 MWCO). The resulting regenerated silk fibroin solution was centrifuged twice at 7799 rcf for 15 min at 4 °C to remove undissolved fibers and debris. The final silk fibroin concentration, referred to as “silk” throughout, was determined via gravimetric analysis by drying a known volume of solution and measuring the residual film. Silk concentrations ranged from 7-9% (wt/vol), and the solution was stored at 4 °C for up to four weeks. Silk solutions were diluted to 1.5 % (wt/vol) in MilliQ water and sterilized by filtration through a 0.22 µm Millex GP PES membrane syringe-driven filter unit (Millipore) using 50 ml plastic syringe and 20 ml silk fibroin solution. The sterilized silk solution was concentrated again between 6-7 % (wt/vol) by dialysis using Snakeskin tubing (3500 MWCO).

Silk hydrogels were prepared by diluting the stock silk fibroin solution in PBS, followed by the addition of Ru/SPS (0.5 mM/5 mM) to get 2% (wt/vol) in final silk concentration. The hydrogel precursor was exposed to a 30 W lamp for 3 mins at room temperature to induce gelation. For NV-loaded hydrogels, NVs suspended in PBS were incorporated into the silk solution to achieve a final silk concentration of 2% (v/v).

### 2.3 Nanovesicle generation and purification

Nanovesicles (NVs) were produced as previously described [54]. In brief, adherent iPSCs were washed twice with PBS, lifted using 10 mM EDTA and centrifuged at 500× g for 5 mins. Cell pellets were resuspended separately in 1 mL of PBS and sequentially extruded through 10, 5, and 1 µm polycarbonate membranes (19 mm; Avanti Polar Lipids, 610010), passing through each filter 13 times using a Whatman extruder. The extruded samples were purified via density gradient ultracentrifugation by layering the extruded sample over 10% and 50% OptiPrep™ (Stemcell Technologies), and centrifuged at 100,000× g for 2 h at 4 °C. The centrifuged samples were fractionated into 6 × 500 µL fractions, each diluted in PBS to a final volume of 1.5 mL to wash NVs. The fractions were centrifuged at 100,000× g for 1 h at 4 °C to pellet NVs (TLA-55 rotor; Optima MAX-MP ultracentrifuge). The resulting NVs were resuspended in PBS and stored at −80 °C until further use. Protein yield for NV fractions was assessed by Micro BCA™ Protein Assay Kit (Thermo Fisher Scientific, 23235).

To evaluate methodological reproducibility, NV generation was performed across two independent experimental periods. Initial experiments included 6 biological replicates processed within a single experimental timeframe to assess *intraday* technical consistency. Subsequently, 3 additional biological replicates were generated 9 months later using identical protocols to evaluate *interday* reproducibility and long-term methodological stability. For proteomic profiling controls, both iPSCs (stem cell control) and HUVECs (non-stem cell control) were cultured under standard conditions as described above. Cells were detached using trypsin, centrifuged at 500× g for 5 mins and lysed with 2% (v/v) sodium dodecyl sulphate (SDS) buffer. Three biological replicates of each cell type were prepared for comparative proteomic analysis to establish baseline stem cell and endothelial cell protein profiles relative to the NV samples. This experimental design enabled assessment of consistency in protein yield, particle size distribution, particle concentration, and expression of key stem cell and tissue repair markers.

### 2.4 Scanning electron microscopy analysis of silk hydrogels

The morphology of NV-loaded and non-loaded hydrogels was studied using scanning electron microscopy as described elsewhere [53]. Three hydrogels from each group were sequentially dehydrated in serial ethanol dilutions (50%, 70%, 90%, 100%, 100%, 100%) for 45 s each dilution using a Biowave Processor (BiowavePro+, Pelco). The hydrogels were further dehydrated via critical point drying (Leica EM CPD300) under soft material cycles with 30 CO_2_ exchanges. The dried samples were cut horizontally, mounted onto carbon tape, and coated with platinum (K575X, EMITECH). The samples were imaged at three random areas per sample using FEI Nova NanoSEM 230 at 15 kV, with the probe current of 30 µA.

### 2.5 Mechanical testing of silk hydrogels

For compression testing, silk hydrogels (with and without NVs) were cast in cylindrical molds (6 mm diameter, 2.5 mm height). Six samples per group were incubated in PBS overnight at 4°C before the compression test. All the hydrogels were compressed to 90% of the initial height at a constant crosshead speed (1 mm s^−1^) at 37 °C in PBS with a 0.1 N pre-load using a mechanical testing machine with a 50N load cell (Instron 5543, USA). Stress–strain curves were plotted for the pre-breaking region, and compressive moduli were determined from the linear region of the stress–strain curves (10–20% strain).

### 2.6 Analysis of the secondary protein structure in silk hydrogels

The secondary structure of silk hydrogels with and without NVs were analyzed using attenuated total reflectance – Fourier transform infrared spectroscopy (ATR-FTIR) as described in previous studies [53, 55] Briefly, the absorbance of freshly prepared hydrogels were acquired in the range of 400-4000 cm^-1^ wavelength at a resolution of 4 cm^-1^ with 24 averaged scans using ATR-FTIR spectrophotometer (Alpha II, Bruker). The obtained Amide I spectrum were then deconvoluted before fitting process using OriginPro software (OriginPro, USA). The position of each peak was further validated by double differentiating the amide I envelopes with peak assignments as follows: side chains/Tyrosine (1595–1605 cm^−1^); β-sheets (1610–1635 and 1697–1704 cm^−1^); random coils (1635–1650 cm−1); alpha-helix (1652–1660 cm^−1^), and beta-turns (1663–1696 cm^−1^)[56].

To visualise the autofluorescence of di-tyrosine bonds, the pure silk hydrogels and NV-loaded silk hydrogels were imaged using GelDoc Gel Imaging system (GelDOC EZ Imager, BioRad) with UV tray under ethidium bromide full spectrum fluorescent settings [53, 55, 56].

### 2.7 Heart tissue sourcing and in-situ tissue adherence test

Access to murine heart tissue in this study was conducted in accordance with the ARRIVE guidelines and approval of the Alfred Research Alliance (ARA) Animal Ethics Committee, Vic, Australia (ethics approval #P10645). Wild-type C57BL/6 mice were sourced from AMREP AS Pty Ltd, VIC. Mice were humanely killed by cervical dislocation, and the chest was immediately open to reveal the heart. 2% silk hydrogel applied directly to the mouse epicardium *in situ*, while the heart was still beating. To evaluate adhesion, two layers of 2.5 µL hydrogel were sequentially applied under dark conditions, with each layer exposed to light for 30 s before applying the next. Following the final layer, the hydrogel was cured with continuous 3-min light exposure *in situ*. The heart was then extracted, and hydrogel adhesion to the myocardial tissue was assessed by subjecting the tissue to flowing water.

### 2.8 Release kinetics of NVs from silk hydrogels

For the NV release kinetics study, 250 μL of 2% silk hydrogel containing 250 μg of NVs was crosslinked in a 6-well plate in triplicates. After crosslinking, 500 μL of PBS was added and the hydrogels were incubated at 37°C for 7 days. PBS was collected and stored at −80°C and fully replaced every 24 h for 7 days. At the end of the experiment, all releasates were concentrated using Amicon^®^ Ultra Centrifugal Filter, 30 kDa MWCO (Millipore, UFC5030) and centrifuged for 13 min at 14 000 g.

### 2.9 Nanoparticle Tracking Analysis (NTA)

Particle concentration and size were determined using nanoparticle tracking analysis (NTA, ZetaView, Particle Metrix, PMX-120; 405 nm laser diode) as per previously described [54]. For NV characterization, 1 μg of NVs was diluted in 1 mL of PBS (n=3, biological). For release kinetics analysis, concentrated released NVs were diluted 1:1000 for Day 1-2, and 1:100 for Day 3-7. A total of 11 positions were captured with the following parameters: camera sensitivity: 80, min area: 5, max area: 1000, brightness: 30, min trace length: 15, temperature: 25 °C. Capture was performed at medium video setting, corresponding to 30 frames per position. ZetaView software 8.5.10 was used to analyze acquired data.

### 2.10 Confocal imaging of labeled-NVs in silk hydrogel

NVs were labelled with fluorescent dye DiI (Vybrant™ DiI Cell-Labeling Solution, 1:200 dilution, Invitrogen, Waltham, MA, USA, V22885) at 1 μM concentration for 15 min at RT as described [54]. Labelled NVs and dye control without NVs were centrifuged at 100,000× g for 1 h at 4°C on 10% and 50% OptiPrep™ cushions (Stemcell Technologies), and pellets were resuspended in PBS and stored at −80°C until used. Labelled NVs and control dye were then encapsulated in 2% silk hydrogels. Hydrogels were prepared the same way as for release kinetics (See method 2.7). After 28 days of incubation at 37°C, a portion of the hydrogel was removed using a surgical blade and placed in an ibidi® µ-Slide chamber in PBS prior to imaging by Nikon A1R confocal microscope equipped with resonant scanner, using a 20× WI (1.2 NA); (Nikon, Tokyo, Japan).

### 2.11 TGF-mediated fibroblast activation assay

Primary human cardiac ventricular fibroblasts (hVCFs) were plated (30,000 cells/cm^2^) on gelatine-coated 24-well plates and allowed to attach overnight. Cells were serum starved for 48 h in DMEM/F12 basal medium to induce quiescence, then activated with 10 ng/mL of TGF-β or equivalent volume of PBS (control) for 72 h, as described [57]. To assess anti-fibrotic function, NVs (30 μg/mL) were added simultaneously with TGF-β treatment [54]. After 72 h, cells were processed for either proteomic or microscopy analysis.

For proteomic analysis, cells were washed in PBS and lysed using 2% (v/v) sodium dodecyl sulphate (SDS) containing Pierce™ Universal Nuclease (Thermo Fisher Scientific, 88700) and HALT protease phosphatase inhibitor cocktail (Thermo Fisher Scientific, 78442).

For microscopy analysis, cells were washed with PBS, fixed with 4% formaldehyde for 5 min, and washed three times with PBS. Cells were permeabilized with 0.2% Triton X-100 for 5 min, then blocked with blocking buffer (3% BSA in 0.2% Triton X-100) for 30 min at room temperature (RT). Primary antibody anti-α-smooth muscle actin was incubated (αSMA; Abcam ab7817, 1:50 dilution) for 1 h at RT, washed with PBS and followed by Alexa Fluor 488 goat anti-mouse secondary antibody (Thermo Fisher Scientific A-11001, 1:200 dilution) for 20 min at RT. Nuclei were counterstained with Hoechst 33342 (5 μg/mL; Thermo Fisher Scientific H1399) according to manufacturer’s instructions. Images were acquired using a Nikon ECLIPSE Ji imaging system, capturing full well views. The αSMA-positive area was quantified and normalized by nuclei count.

To assess the anti-fibrotic function of silk-NV composite, 50 μL of 2% silk hydrogels without (empty control) or with low (500 μg/mL hydrogel) or high (700 μg/mL hydrogel) concentration of NVs were cast directly in the upper chamber of a 24 well plate Transwell® with polycarbonate membrane of 8 μm pore size (Corning, CLS3422) and placed above quiescent hVCFs. All treatments were performed in DMEM/F12. After 72h of treatment, cells were washed twice in PBS and lysed in 2% (v/v) sodium dodecyl sulphate (SDS) for proteomics and western blot analysis.

### 2.12 Western blotting

For immunoblotting, cell lysates were mixed 1:1 with loading buffer (4% SDS, 20% glycerol, 0.01% bromophenol blue, 0.125 mM Tris-HCl, pH 6.8, with 1M DTT) and separated on NuPAGE™ 4-12% Bis-Tris gels (Invitrogen, NP0321) at 150 V for 1 h. Proteins were transferred to nitrocellulose membranes using iBlot™ 2.0 Dry Blotting System (Life Technologies) at 20 V for 7 min. Membranes were blocked with skim milk in TPBS (PBS with 0.1% Tween 20) for 1 h at room temperature with agitation, followed by three 5-min TPBS rinses. Primary antibodies against α-sMA (1:1000, Abcam, ab5694) and GAPDH (1:1000, Cell Signaling Technology, D4C6R/97166S) were applied overnight at 4°C in TPBS. After washing, membranes were incubated with secondary antibodies (1:20,000, IRDye 800CW goat anti-mouse and IRDye 680RD goat anti-rabbit; LI-COR Biosciences, 926-68071 and 926-32210) for 1 h at room temperature. Following three TPBS washes, membranes were imaged using Odyssey Infrared Imaging System (LI-COR Biosciences) at 700 and 800 nm. Expression intensity was assessed using ImageJ.

### 2.13 Murine Ischemia Reperfusion Injury model and patch application

All animal procedures were conducted in accordance with guidelines approved by the Alfred Research Alliance Animal Ethics Committee (Vic, Australia; ethics approval P10645). Wild-type male C57BL/6 mice (10–12 weeks old, 20–30 g) were sourced from AMREP AS Pty Ltd and housed under standard conditions with ad libitum access to food and water. Mice were randomly assigned to sham (n = 6), ischemia–reperfusion injury (IRI, n = 6), IRI + empty silk patch (n = 8), or IRI + silkNV (n = 7) groups.

Mice were anesthetized by intraperitoneal injection of ketamine/xylazine/atropine (80–100 mg/kg, 16–20 mg/kg, and 0.96–1.2 mg/kg, respectively). Following induction of anesthesia, animals were intubated and mechanically ventilated. A left thoracotomy was performed to expose the heart, and ischemia–reperfusion injury was induced by temporary ligation of the left anterior descending (LAD) coronary artery using a non-absorbable suture. After 60 min of ischemia, the ligature was released to allow reperfusion. Sham animals underwent identical surgical procedures without LAD ligation.

At the time of reperfusion, mice received either no treatment (IRI), a silk hydrogel patch alone (IRI + empty patch), or an NV-loaded silk hydrogel patch (IRI + silkNV) applied directly to the epicardial surface overlying the infarct region. Silk hydrogel (2% w/v; 5 µL per heart) containing ruthenium (0.5 mM) and sodium persulfate (5 mM) was photocrosslinked in situ using visible light to form the patch. For the silkNV group, iPSC-derived nanovesicles were incorporated into the hydrogel at 1 g/mL prior to application.

Animals that failed to recover as expected or exhibited signs of abnormal pain or distress were humanely euthanized in accordance with ethical guidelines. The overall surgical mortality rate for this model was approximately 25%. Exclusion criteria for downstream analysis included mice exhibiting a left ventricular ejection fraction greater than 40% and an area at risk less than 10%, as determined by echocardiography 24 h post-surgery, which were deemed surgical failures and excluded from the study. Animals were monitored closely throughout recovery. Hearts were harvested at 28 days post-injury for downstream analyses.

### 2.14 Echocardiographic assessment of cardiac function

Echocardiography was performed at baseline, 1 day and 28 days post IRI using a Vevo2100 ultrasound system (VisualSonics, Toronto, ON, Canada) equipped with an MS550D transducer to assess cardiac function. Mice were anesthetized using inhaled isoflurane (3–4.5% for induction and 1–2% for maintenance in oxygen) and positioned supine on a heated imaging platform to maintain body temperature throughout the procedure. Eye lubrication was applied to prevent corneal drying. B-mode loops of the left parasternal long-axis view of the heart were captured. LV volumes at end diastole (LVEDV) and end systole (LVESV) were obtained, and LV ejection fraction (EF) was calculated. All image acquisition and analyses were performed by operators blinded to treatment groups.

### 2.15 Histology

At 28 days post-surgery, hearts were harvested and fixed in 10% neutral buffered formalin (NBF). Fixed tissues were processed and paraffin-embedded by the AMREP Monash Histology Platform. Transverse cardiac sections (6 µm thickness) were prepared and stained with either Sirius Red to assess collagen deposition and fibrotic remodeling or hematoxylin-eosin (H&E) staining for tissue morphology and toxicity of the silk patch. Whole-section images were acquired by brightfield microscopy. Fibrotic area was quantified as a percentage of total left ventricular area using QuPath, and minimum left ventricular wall thickness was measured at the thinnest region of the infarct. All analyses were performed in a blinded manner.

### 2.16 Heart tissue homogenization and lysis

Left ventricle Apex and Base tissues allocated for proteomic analysis were lysed in 1 mL of lysis buffer (50 mM Tris–HCl, 2% SDS, 10 mM EDTA in PBS, pH 8) supplemented with HALT protease and phosphatase inhibitor (Thermo Fisher Scientific). For tissue homogenization, three 3.2-mm stainless steel beads were added to each tube and maintained on ice. Samples were then processed using Bullet Blender 24 Gold (Next Advance) for 1 min (setting 8; 30-s pulse), with tubes placed on ice between pulses. Homogenates were further disrupted by probe sonication using a microtip (23 amplitude, 10 s) at 4 °C. Lysates were centrifuged at 17,000 × g for 20 min to pellet insoluble debris, and supernatants were collected for protein quantification using a microBCA assay (Thermo Fisher Scientific) according to the manufacturer’s instructions. Protein lysates were stored at −80 °C until downstream analysis.

### 2.17 Mass spectrometry-based proteomics

For NV characterization, iPSC-derived nanovesicles (iPSC-NVs; n = 9, biological replicates including inter- and intra-batch) were lysed in 2% (v/v) sodium dodecyl sulfate (SDS) containing 50 mM HEPES (pH 8.0) and HALT protease and phosphatase inhibitors (78442, Thermo Fisher Scientific), alongside cell controls including iPSCs (n = 4, biological) and HUVECs (n = 3, biological).

For in vitro anti-fibrotic assays, primary human ventricular cardiac fibroblasts (hVCFs) were lysed at the assay endpoint (see section 2.9) following TGF-β activation (n = 3) with or without treatment with silk-NVs (n = 4), empty silk (n = 4), or NVs alone (n = 4).

For in vivo proteomic remodelling assessment, mouse left ventricular cardiac tissues (Apex and Base regions) were collected 28 days post-injury from sham (n = 6), IRI (n = 6), IRI + empty patch (n = 8), and IRI + silkNV (n = 7) groups. Tissue lysates were prepared as described in Section 2.14.

For all samples, protein concentrations were determined using BCA assay, and 5 µg of protein were processed for downstream analysis. Samples were reduced and alkylated with 10□mM DTT and 20 mM IAA and as described [58] using solid-phase-enhanced sample preparation. Samples were digested using Trypsin and Lys-C (1:50 and 1:100 enzyme-to-protein ratio, respectively) overnight at 37°C. Samples were acidified after digestion to final concentration of 2% formic acid before vacuum lyophilisation. Samples were reconstituted in 10□µ□ of 0.07%(v/v) trifluoroacetic acid in LC-MS grade water and peptide concentration determined using Pierce™ Quantitative Fluorometric Peptide Assay (Thermo Fisher Scientific, 23290). LC-MS data acquisition was performed on Q Exactive HF-X benchtop Orbitrap mass spectrometer coupled with UltiMate™ NCS-3500RS nano-HPLC and operated with Xcalibur software as described [59, 60]. MS-based proteomics data is deposited to the ProteomeXchange Consortium via the MassIVE partner repository and available via MassIVE with identifier MSV MSV000098711.

DIA-MS spectra were processed using DIA-NN software (v1.9) [61] as reported [59, 62]. The DIA-MS spectra were searched against human proteome database (UP000005640, #83,401). Perseus (v1.6.13) was applied for data processing and analysis, with scatter plots/bar charts generated using GraphPad Prism (v9.2.0) or Microsoft Excel. The mass spectra were analyzed using default settings with a false discovery rate (FDR) of 1% for precursor identifications. For downstream bioinformatics analysis, we applied a cut-off (protein identification inclusion) of at least 70% valid values in total in at least one group. Protein intensities were transformed into log2 and normalized using quantile normalization. Proteins were subjected to PCA and unpaired student’s t-test or Welch’s t-test with missing values imputed from normal distribution (width 0.3, downshift 1.8). g:Profiler [63] and Reactome [64] pathway databases were utilized for Gene Ontology functional enrichment and network/pathway analysis, significance P<0.05. Venn diagram was generated using a web-based tool (http://www.interactivenn.net/) [65]. To identify surface annotated proteins in the NV proteome we performed surface annotation analysis using in silico surfaceome SURFY [66], cell surface protein atlas CSPA [67], and cell surface protein annotation using UniProt [68]. We further analysed NV proteome for identified peptide sequence comprising tyrosine (Y) resides. Functional protein network analysis was performed using STRING 12.0 (https://string-db.org/)[69].

### 2.18 Statistics

Data are expressed as mean ± standard error of the mean (SEM). Significance of the differences was evaluated using Student’s or Welch’s t-test, one-way or two-way ANOVA. P < 0.05 is considered statistically significant.

## 3. RESULTS

### 3.1. Consistent generation of NVs with therapeutic cargo

Nanovesicles (NVs) were generated through a rapid cell extrusion workflow using RM3.5 GT-Td Tomato human iPSCs [70] as donor cell source (**Fig. 1A**). iPSC cell suspensions were serially extruded trough membranes of decreasing pore sizes, followed by density gradient ultracentrifugation for purification and isolation of the NV fraction from soluble, non-membranous factors (**Fig. 1B-C**). To assess yield and intra-batch consistency of NV generation, we performed 18 extrusions from independent biological replicates, pooling these into six NV preparations (referred to as batch 1) (**Fig. 1B**) as well as 3 additional extrusions performed on separate days to assess inter-batch consistency (referred as batch 2). Our NV generation workflow resulted in an NV yield of ∼25 µg/million cells (**Fig. 1D**). Single particle tracking analysis revealed particle size/diameter (range 50-300 nm, mean: ∼110 nm, n=3, biological) and particle concentration (∼1.4 × 10¹¹ particles/mL) (**Fig. 1E, Supplementary Fig. S1A**).

**Figure 1.**
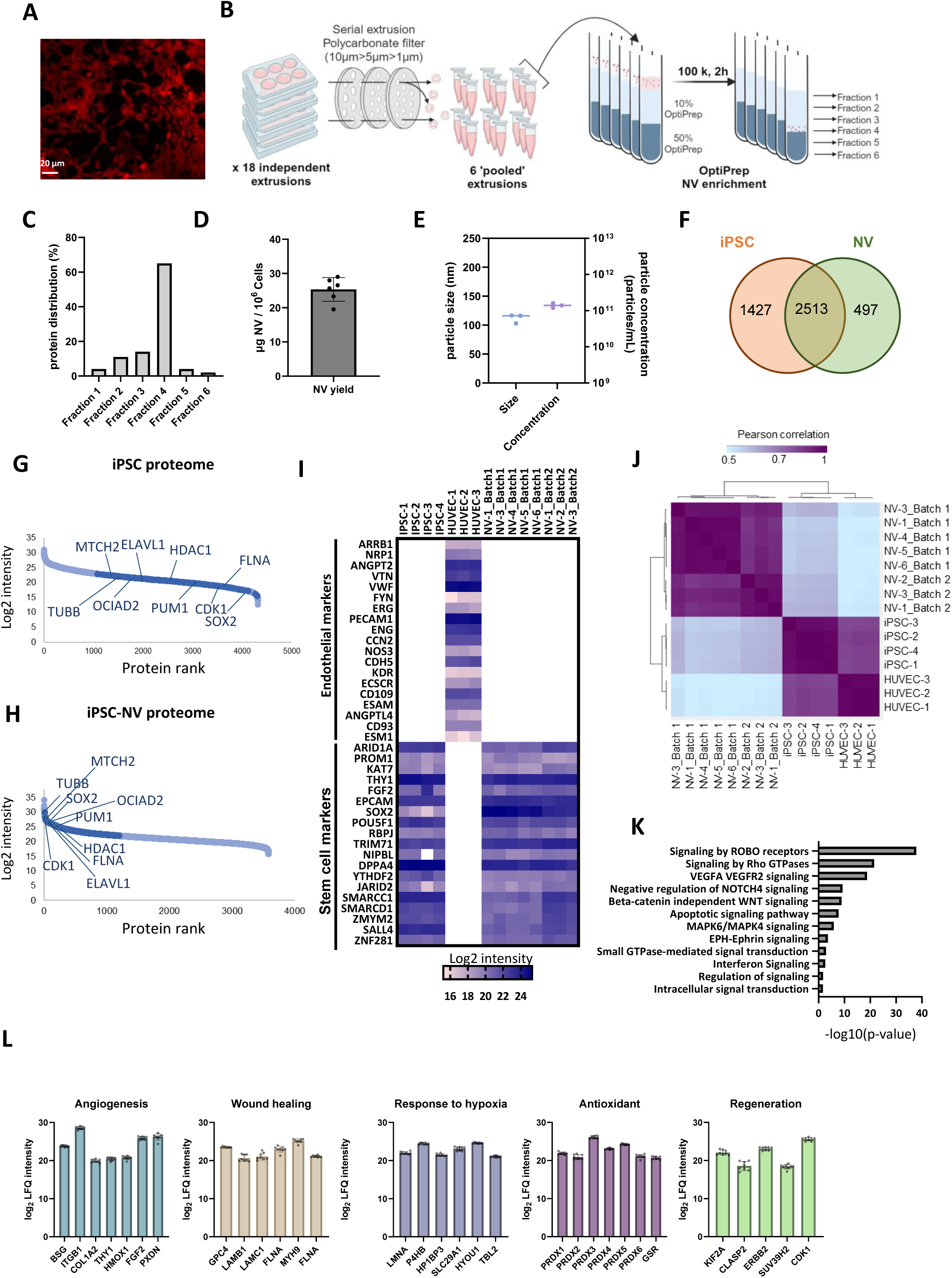
Generation and proteomic profiling of iPSC-derived nanovesicles. (**A**) Fluorescence microscopy image of RM3.GT-TdTomato iPSCs (20× magnification, Olympus IX71 microscope). (**B**) Schematic overview of NV generation (18 independent preparations, across different batch generations) from iPSCs by serial extrusion and OptiPrep density-cushion ultracentrifugation, yielding fractions 1–6. (**C**) Protein concentration distribution across the density gradient fractions, assessed by micro-BCA assay (expressed as percentage of total protein); fraction 4 identified as NV-enriched (mean, 6 biological). (**D**) Protein yield from NV-enriched fraction 4, quantified by micro-BCA and expressed as µg of NVs per million cells (n = 6, mean ± SD, biological). (**E**) Particle size distribution and concentration of NVs, determined by nanoparticle tracking analysis (NTA) (n = 3, biological). (**F**) Venn diagram showing shared and unique proteins between iPSC and NV proteomes. (**G-H**) Abundance distribution (waterfall plots, LFQ intensity, log_10_) of proteins identified in iPSCs (**G**) and derived NVs (**H**). Highlighted in darker blue are proteins enriched in NVs in comparison to iPSCs. (**I**) Heatmap of selected endothelial and stem cell marker expression in iPSC, HUVEC, and NV proteomes. (**J**) Pearson correlation matrix of NV protein profiles across extrusion replicates from batch 1 (n = 5, biological) and batch 2 (n = 3, biological), as well as iPSC (n = 4, biological) and endothelial (HUVEC) control (n = 3, biological). (**K**) Gene Ontology (GO) enrichment analysis of biological processes in the NV proteome showing enrichment in signalling proteins. (**L**) Bar graph of logL LFQ intensity values for selected proteins found in 8 out of 8 NV replicates and associated with angiogenesis, wound healing, antioxidant activity, response to hypoxia, and tissue regeneration (UniProt annotations).

To understand the proteome composition of iPSC-NVs relative to their cell source and the impact of cell type compared to a non-stem cell source, we performed mass spectrometry-based proteomic analysis, similar to our previous studies [54]. For NV proteome composition, we performed proteomic analysis on iPSC-derived NVs (n=6 for batch 1 and n=3 for batch 2), in comparison to both source iPSCs (n=4, biological) and HUVECs (n=3, biological) (**Supplementary Fig. S1B, Table S1**). Proteomic profiling identified 3940 and 3010 proteins in iPSC and iPSC-NVs, respectively, indicating a high similarity in their proteomes (**Fig. 1F, Table S1**). Differences in proteome landscape reveal enrichment of membrane and membrane-associated organelles in NV, relative to the complexity of donor cell proteome (**Supplementary Fig S1.C, Table S2**). We further highlight 183 surface annotated proteins in NV proteome, annotated using a combination of Cell Surface Atlas annotation databases [66–68] (**Supplementary Fig S1.E, Table S1**). To gain insights into potential interaction of NVs with silk crosslinking and tyrosine-mediated interface, we further identify 57/183 surface proteins in NV proteome as containing a tyrosine residue **(Table S3).**

Principal component analysis showed that iPSC-NVs and iPSCs proteomes were distinct from HUVEC proteome (**Supplementary Fig. S1C**). The NV proteome displayed a dynamic range of protein abundance spanning over 6 orders of magnitude (based on LFQ intensities), with proteins ranked according to their abundance relative to iPSC cell proteome (**Fig. 1G-H, Table S4**). NVs were enriched in pluripotent markers SOX2, POU5F1/OCT4, and TRIM71 (**Fig. 1H, Table S1**), aligning with their iPSC origin. NVs also did not express non-pluripotent and linage-specific markers such as endothelial markers, confirmed by comparison to endothelial and stem cell-enriched proteins from Human Protein Atlas (HPA) [71] (**Fig. 1I, Table S1**). Based on biophysical (protein yield, particle size, particle concentration) (**Fig. 1D-E**) and proteome composition (**Fig. 1J**), we demonstrate that NV generation is highly consistent. When comparing batch consistency based on the total NV proteome, Pearson correlation analysis demonstrated high reproducibility both between batches (r>0.85) and within batches (r>0.95) (**Fig. 1J**).

Ontology analysis of iPSC-NV proteome revealed significant enrichment in complex signaling networks, including Rho GTPase signaling, VEGFA/VEGR signaling and EPH-Ephrin signaling (**Fig. 1K, Table S5**). Further, based on ranked abundance in NVs and consistent expression across biological replicates, we highlight key biological processes associated with NV cargo, including proteins associated with angiogenesis (FGF2, HMOX1, BSG), wound healing (LAMB1, FLNA, GPC4), antioxidant activity (PRDX1–6, GSR), tissue regeneration (KIF2A, CDK1, ERBB2) and hypoxic response (HYOU1, P4HB) **(Fig. 1L, Table S1)**. We further identify 63 proteins, detected in at least 7 out of 9 replicates, as markers enriched in heart tissue (i.e., CDH2, NDUFS1, ACO2, MYL12A), based on the Human Protein Atlas global tissue dataset (**Table S1**), highlighting functional and therapeutic relevance of iPSC-NVs as therapeutics for heart repair.

### 3.2. NVs have anti-fibrotic activity and remodel myofibroblasts proteome

Fibroblast activation is a hallmark of cardiac pathological remodeling contributing to heart failure. In this context, TGFβ is a driver of maladaptive cardiac fibroblast biology, promoting myofibroblast differentiation and ECM deposition [72]. We induced a pathological phenotype in human ventricular fibroblasts by treating cells with TGFβ for 72 h and measured α-SMA expression as a marker of myofibroblasts (**Fig. 2A**). We then assessed whether co-treatment with NVs (30 µg/mL) could rescue fibroblast activation (**Fig. 2A**). Using western blot analysis and immunohistochemistry, we found that NV treatment significantly reduced α-SMA expression (by 2-fold, *p<0.0005*), attenuating α-SMA expression relevant to non-activated fibroblasts (**Fig. 2B-C and Supplementary Fig. S2A-B**).

**Figure 2.**
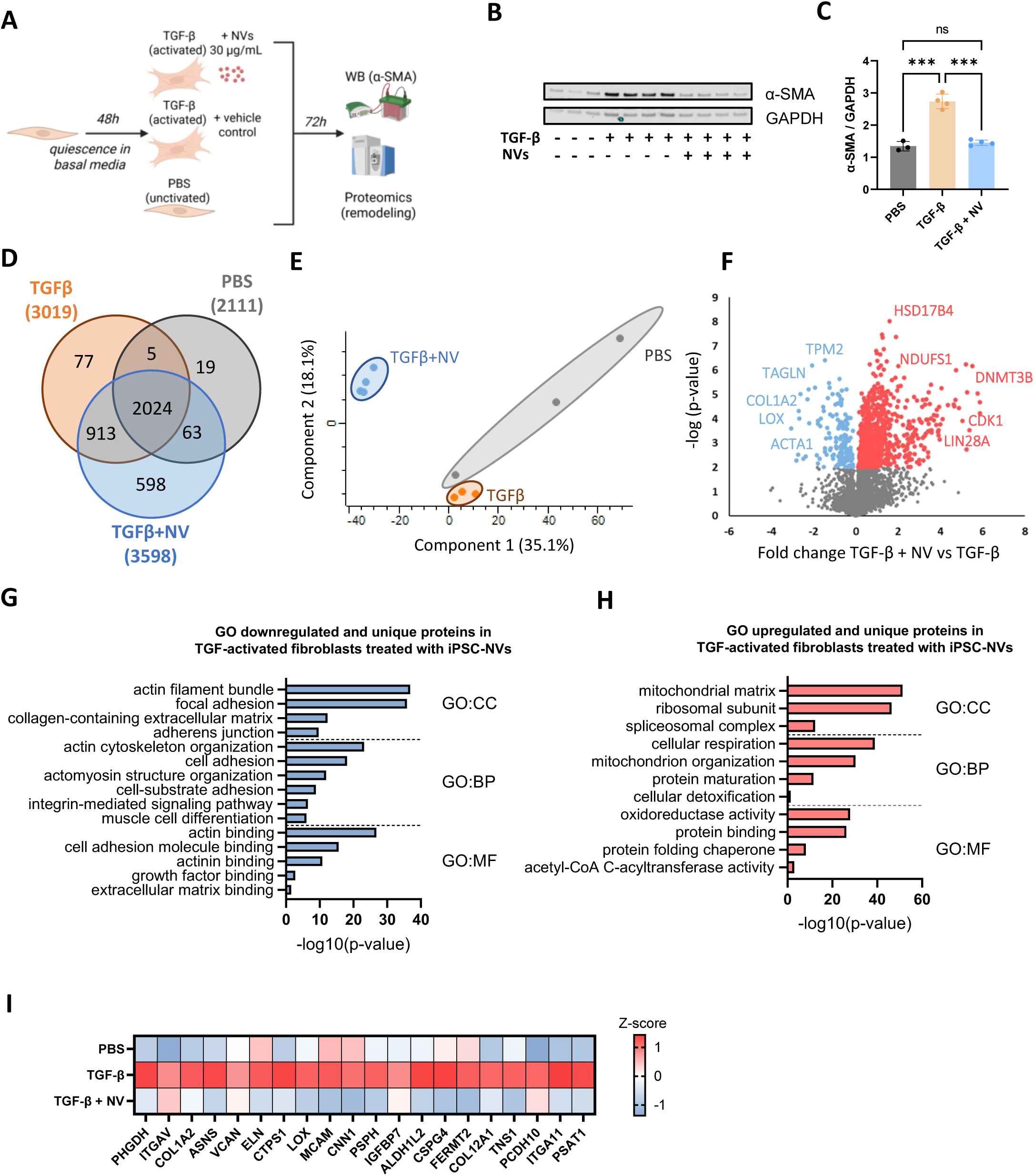
Function and remodeling of TGF-treated cardiac fibroblasts by iPSC-derived NVs. (**A**) Experimental workflow of TGFβ-mediated human ventricular cardiac fibroblast (hVCFs) activation and NV treatment assay. hCVFs were cultured under PBS (inactivated), TGFβ (activated), or TGFβ+NV (30□µg/mL) conditions. After 72 hours, α-SMA expression quantified via Western Blot analysis and fibrotic remodelling by proteomic analysis. (**B**) Western blot of α-SMA in PBS, TGFβ, and TGFβ+NV-treated hVCFs (n□=□3, PBS, n□=□4, all; biological). (**C**) Quantification of α-SMA expression normalized to GAPDH (mean□±□SD, two-way ANOVA with Tukey’s multiple comparison). (**D**) Venn diagram of identified proteins across PBS, TGFβ, and TGFβ+NV conditions. (**E**) Principal Component Analysis (PCA) of hVCFs proteomes in treatment condition PBS, TGFβ, and TGFβ+NV. (**F**) Volcano plot comparing TGFβ+NV vs TGFβ treated hVCFs (t-test pL<□0.05; red—upregulated, blue—downregulated in TGFβ+NV). (**G-H**) Selected Gene Ontology (GO) terms enriched in (**G**) downregulated proteins in TGFβ+NV vs TGFβ (and those unique to TGFβ) (**H**) and upregulated proteins in TGFβ+NV vs TGFβ (and those unique to TGFβ+NV); CC: Cellular Component, BP: Biological Process, MF: Molecular Function. (**I**) Heatmap of z-score LFQ intensity of known fibrotic markers in PBS, TGFβ, and TGFβ+NV conditions.

Next, we sought to understand the biological response in cardiac myofibroblasts following direct NV treatment by analyzing their reprogrammed cell proteome (**Figure 2D, Sup Table S6**). Principal component analysis revealed that vehicle (PBS), TGFβ, and TGFβ+NV treated cell proteomes clustered independently (**Fig. 2E**). Pairwise comparative analysis (TGFβ vs PBS) revealed 439 significantly dysregulated proteins (*Student’s T-test*, *p* < 0.05, 214 upregulated, 225 downregulated) in response to TGFβ including COL1A2, ITGA11, LOX, and ACOT1 (**Supplementary Fig. S2C, Table S6**). We further highlight enrichment in networks associated with ECM remodeling (COL1A1, THBS1, LOX), actin cytoskeleton organization (FLNB, PARVB), and cell adhesion (MCAM, ITGAV, FAP), alongside downregulation of mitochondrion organization (NDUFS1, AFG3L2, MCU), reflective of a fibrotic phenotype (**Supplementary Fig. S2D-E. Table S7 and S8**).

Pairwise comparative analysis of NV treatment response, relative to TGFβ, revealed 1725 significantly dysregulated proteins (*Student’s T-test*, *p* < 0.05, 1574 upregulated, 151 downregulated), including COL1A2 and LOX (**Fig. 2F, Table S6**). In TGFβ+NV, Gene Ontology (GO) analysis of downregulated proteins (*Student’s T-test p* < 0.05 or uniquely identified in TGFβ) revealed protein networks associated with collagen-containing ECM (LTBP1, VCAN, COL1A1/2, FN1) and actomyosin structure organization (FLNC, MYH9/10, ACTA1, CNN1) (**Fig. 2G, Table S9**), alongside upregulation of metabolic processes, mitochondrial organization, and oxidoreductase activity (**Fig. 2H, Table S10**). We highlight NV-mediated downregulation of matricellular proteins (VCAN and ELN), metabolism proteins (PHGDH, ASNS, CTPS1) and adhesion proteins (ITGAV, ITGA11, MCAM, FERMT2), relative to TGFβ alone (**Fig. 2I**).

Thus, our functional and proteomic data support NV’s anti-fibrotic function in activated cardiac fibroblasts.

### 3.3. Developing porous silk biomaterial for NV encapsulation

To achieve sustained and localized delivery of NVs, we developed a silk-based hydrogel delivery platform by encapsulating NVs within silk fibroin matrix via di-tyrosine photo-crosslinking [52] (**Fig. 3A**). This visible light-mediated reaction, catalyzed by ruthenium (Ru) and sodium persulfate (SPS), induces covalent di-tyrosine bond formation between tyrosine residues[53]. NV-loaded silk hydrogels displayed a significantly smaller diameter than empty hydrogels, with average diameters of 5.1 mm and 5.3 mm, respectively (**Fig. 3B–C, Table S11**). This reduction in size suggests that the incorporation of NVs led to a denser crosslinked network. Scanning electron microscopy (SEM) revealed a porous, heterogeneous fibrillar architecture in both hydrogels, arising from the self-assembly of silk into nanoscale protein spheres that coalesce into a fibrous network (**Fig. 3D**). NV incorporation did not alter the overall pore structure, but NVs were observed to associate with silk fibrils and be retained within the network pores, indicating integration within the hydrogel matrix (**Fig. 3D**). Degradation rate assessed by wet hydrogels mass loss showed that both hydrogels degraded at a similar rate for the first 7 days, followed by a significant decrease in degradation rate in the NV-loaded silk at 12 and 20 days compared to empty silk control **(Supplementary Fig. S3 A)**. Mechanical testing demonstrated that NV-loaded hydrogels exhibited an increased compressive modulus and enhanced toughness relative to empty hydrogels (Fig. 3E–G, Table S12). Given the observed increase in compressive modulus following NV incorporation, we next investigated whether these changes were associated with alterations in the secondary structure of silk fibroin using attenuated total reflectance Fourier transform infrared (ATR-FTIR). Indeed, NV incorporation influenced the secondary structure of silk fibroin, with NV-loaded hydrogels exhibiting increased β-sheet content and a reduction in random coil structures (**Fig. 3 H-I)**. Notably, these molecular changes occurred without observable differences in hydrogel porosity by SEM (**Fig. 3D)**. Together, these findings suggest that the improved mechanical properties of NV-loaded hydrogels arise from an increased β-sheet-mediated physical crosslinking. Finally, autofluorescence, indicative of di-tyrosine crosslinking, was higher in empty silk hydrogels than in NV-loaded hydrogels. This difference may be influenced by the increased opacity of NV-loaded samples, which can reduce light transmission and apparent signal intensity rather than a decrease in di-tyrosine crosslinking density (**Supplementary Fig. S3 B)**.

**Figure 3.**
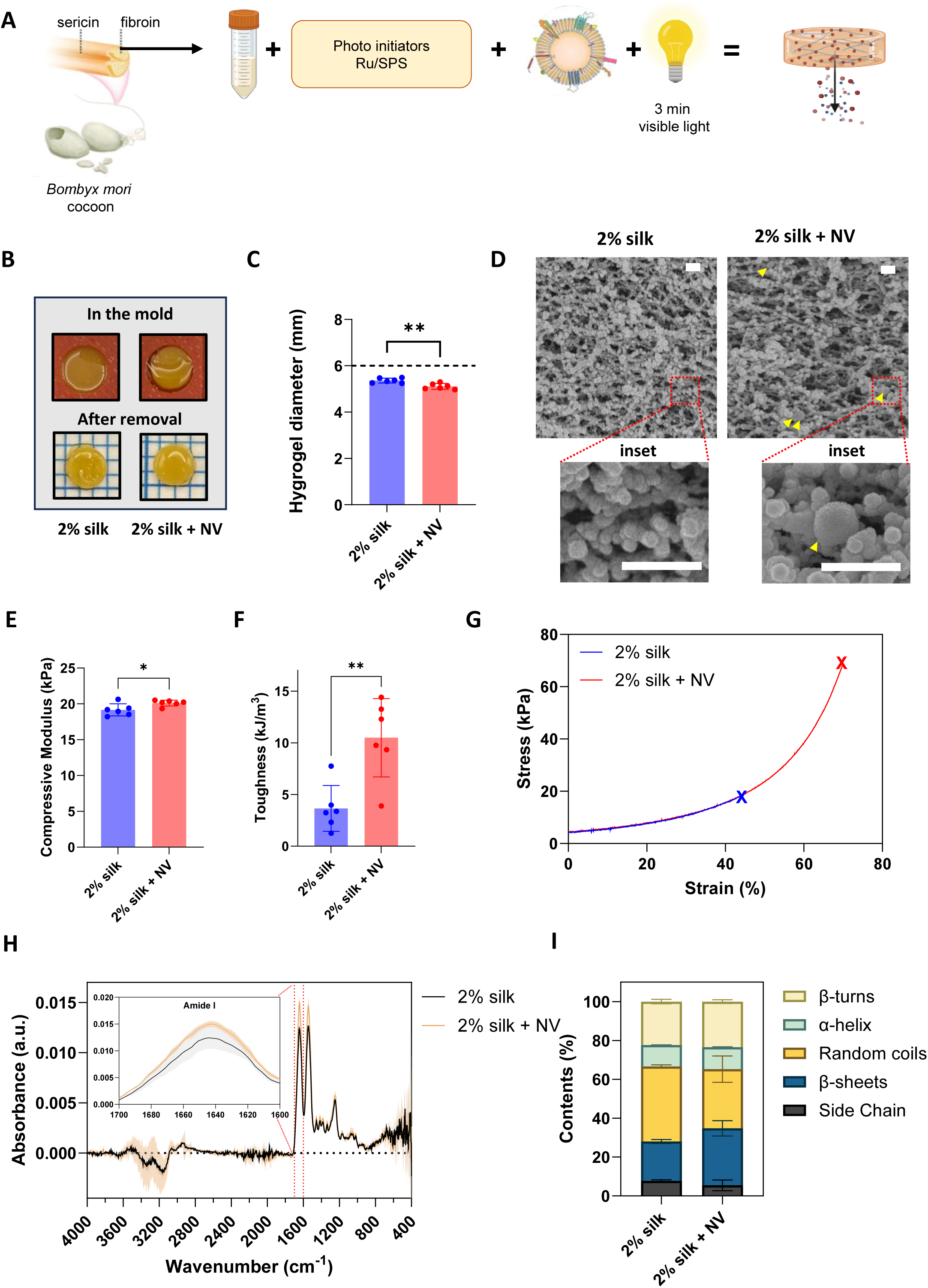
Preparation and characterization of NV-loaded silk hydrogel. (**A**) Schematic of experimental workflow for NV encapsulation in silk fibroin hydrogel using Ru/SPS photo-initiators and visible light for crosslinking. Concentration: 35 ug of NV per 50 uL hydrogel patch (700ug NV/mL hydrogel). (**B**) Representative photographs of 2% silk hydrogels without (left) and with NVs (right) postLcrosslinking and after being removed from the mold. (**C**) Diameter of the formed hydrogels as measured using Image J (n=6, mean□±□SD, t-test). The black dashed line indicates the initial mold size 6mm. (**D**) Representative SEM images of silk hydrogels without (left) and with NVs (right). White scale bar is 500 nm. Yellow arrows indicate presence of NVs. (**E-G**) Shear modulus analysis of silk hydrogels (n=5) showing the compressive mechanical testing (**E**), toughness (**F**), stress-strain curves(**G**). (**H**) Attenuated total reflectance – Fourier transform infrared (ATR-FTIR) spectrum absorbance of silk hydrogels without (2% silk) and with NVs (2% silk + NV) identifying Amide I region (n=5 hydrogels/group) **(I)** Quantitative analysis of ATR-FTIR of Amide I region showing β-turn, α-helix, random coils, β-sheet and side chaines content in silk hydrogels without (2% silk) and with NVs (2% silk + NV).

To demonstrate proof-of-concept attachment and adhesive properties directly to heart tissue, we evaluated direct application to pericardium (external heart tissue). Here, *in-situ* adhesion of the photo-crosslinked silk hydrogel on beating murine hearts was performed (**Supplementary Fig. S3 C, video 1**). NV-loaded hydrogels remained firmly attached to the myocardium (30 min post cross-linking), resisting external force by flushing running water (**Supplementary Fig. S3 D, video 2**). Thus, we demonstrate that iPSC-derived NVs can be successfully incorporated into silk fibroin hydrogels using a di-tyrosine photo-crosslinking strategy, contributing to silk mechanical integrity and suitable for injection-free administration to the epicardium.

### 3.4. Sustained release of functional NVs from silk fibroin hydrogels

We next evaluated whether NVs released from silk fibroin hydrogels retained their biophysical integrity and functional properties. We hypothesize that NVs are partially adhered to the silk hydrogel network, enabling the biomaterial to function as a delivery system and an immobilization platform for NVs. We demonstrate NV release kinetics over a 7-day period, with significant release of NVs in the first 2-3 days, followed by sustained release over 7 days (**Fig. 4A, Supplementary Fig. S4A-G**). Empty silk hydrogels also resulted in particle release in the first 2 days, likely due to inherent particle release (**Fig. 4A, Supplementary Fig. S4A-G**). We further show that NV biophysical characteristics and bioactivity was retained following their incorporation and release from silk hydrogels, including particle size, range, and diameter (**Fig. 4B**, **Supplementary Fig. S4H-G**). Notably, DiI-labelled NVs were detected in the hydrogel after 28 days *in vitro* (**Fig. 4C**). Additionally, NV-loaded hydrogels significantly reduced α-SMA expression over 72 h in a dose-dependent manner compared to empty hydrogels (**Fig. 4D-F**). We show that released NVs not only preserved their biological activity but significantly contributed to the functional properties of this biomaterial platform in a model of anti-fibrotic activity on primary cardiac fibroblasts.

**Figure 4.**
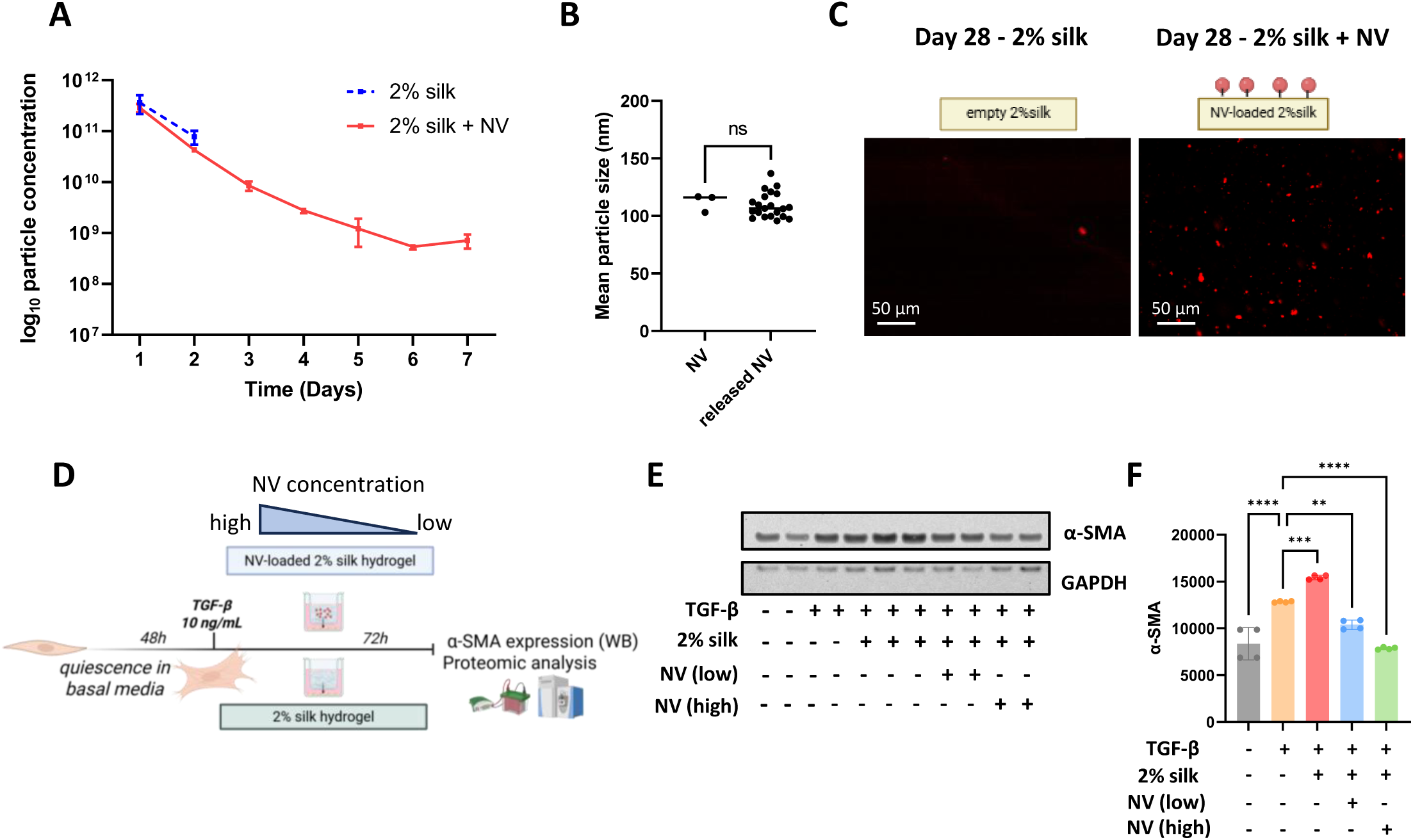
Sustained release of intact and functional iPSC-derived NVs from silk fibroin hydrogels. (**A**) Release kinetics of NVs from 2% silk hydrogels incubated at 37°C in PBS over 7 days, quantified by nanoparticle tracking analysis (NTA) and expressed as log□□ particle concentration (n□=□3, biological). Grey dotted line represents particles released from empty silk hydrogel control, for which particles were below the detection threshold after Day 2. (**B**) Average particle size of control NVs vs pooled released NVs (days 1–7) as assessed by NTA (n=3 per timepoints, t-test, ns: not significant). (**C**) Confocal image of DiI-labelled NVs encapsulated in 2% silk hydrogel or empty silk control after 28 days (scale bar□=□50□µm). (**D**) Experimental workflow of TGFβ-mediated cardiac fibroblast activation using a transwell system with PBS (inactivated), TGFβ (activated), TGFβ+empty silk, or TGFβ+NV-loaded silk (low: 500□µg/mL, high: 700□µg/mL); α-SMA expression assessed at 72□hours by Western blot. (**E**) Western blot image of α-SMA and GAPDH expression across treatment groups (n□=□4). (**F**) Quantification of α-SMA expression (mean□±□SD, two-way ANOVA with Tukey’s multiple comparison).

### 3.5. NV-biomaterial platform reprograms cardiac fibroblast proteome

To understand the interaction between silk and loaded NVs on cell signaling relative to direct NV application, we performed proteomic analysis on myofibroblasts treated for 72 h with TGF-β and either NV-loaded or empty silk hydrogels placed in a transwell system **(Fig. 4D)**. Proteomic profiling demonstrated NV-associated modulation of fibrosis-related pathways, with comparable effects between direct NVs (non-released) and biomaterial-released NVs. Heatmap expression analysis (p<0.05, one-way ANOVA) revealed a subset of proteins downregulated exclusively in released NV group compared to TGFβ alone or empty silk hydrogels (**Supplementary Fig. S5A**). GO analysis of this subset revealed their association with focal adhesion (CNN1/3, NEXN, CSPG4), actin filament bundle (FLNA, FERMT2, CALD1), and smooth muscle contraction (MYLK, TPM1/4, VCL) (**Supplementary Fig. S5B**).

To assess the effects of released NVs on the proteomic remodeling of cardiac fibroblasts, we performed pairwise comparison between TGF-β+silk_NV and TGF-β+silk, identifying 221 dysregulated proteins (*Student’s T-test* p < 0.05), including 122 upregulated and 99 downregulated proteins (**Fig. 5A, Table S13**). Downregulated proteins were associated with actin cytoskeleton organization (LMOD1,CKAP5, FLNA), cell migration (FERMT2, PALLD, TUBA1A), and integrin-mediated signaling pathway (ITGA1, ITGA11, ZYX) (**Fig. 5B, Table S14**), while upregulated proteins were enriched in networks associated with glycolysis (ALDOA, HK2, ALDH7A1), protein folding chaperones (PFDN2, CCT3/8, TOR1A), and protein transport activity (TOMM70, SEC63, TMED10) (**Supplementary Fig. S5C, Table S15**).

**Figure 5.**
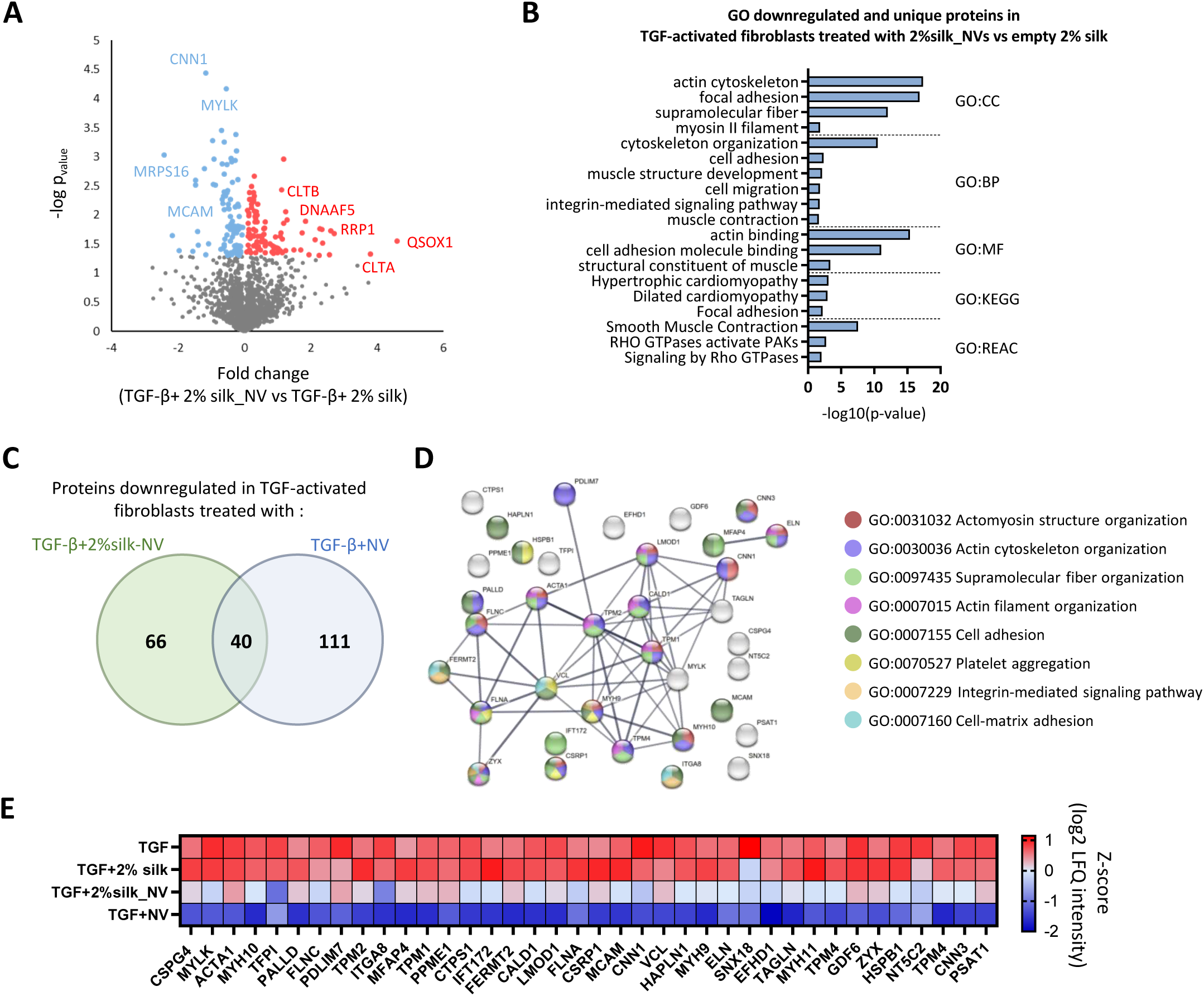
Proteomic reprogramming of cardiac fibroblasts by silk-NV composites. (**A**) Volcano plot comparing TGFβ+2% silk_NV vs TGFβ+2% silk (t-test p□<□0.05; red—upregulated, blue—downregulated in TGFβ+2% silk_NV). (**B**) Selected Gene Ontology (GO) terms enriched in downregulated proteins in the NV-loaded hydrogel group (TGFβ+2% silk_NV) compared with the empty silk hydrogel group (TGFβ+2% silk), or unique to the TGFβ + 2% silk; CC: Cellular Component, BP: Biological Process, MF: Molecular Function, KEGG: Kyoto Encyclopedia of Genes and Genomes. (**C**) Venn diagram of downregulated proteins in both TGFβ+NV and TGFβ+2% silk_NV groups compared to TGFβ, showing 40 shared proteins. (**D**) STRING network analysis of the 40 shared downregulated proteins with associated GO term annotations. (**E**) Heatmap of z-score LFQ intensity for fibrotic markers across TGFβ, TGFβ+2% silk, TGFβ+NV, and TGFβ+2% silk_NV conditions.

To compare proteome remodeling associated with fibrosis regulation following direct NV treatment or NV released from the silk biomaterial, we identified 40 proteins commonly downregulated relative to TGFβ (**Fig. 5C**). Functional network analysis highlighted key protein clusters related to integrin-mediated signaling (ITGA8, FERMT2), actomyosin structure organization (ACTA1, ELN, FLNC), and cell adhesion (HAPLN1, ZYX, PALLD, MCAM) (**Fig 5D**). Ranked-based protein abundance revealed how specific proteins in activated human primary fibroblasts were selectively regulated by NVs (both direct and released from biomaterials) to support their anti-fibrotic function (**Fig. 5E**).

To further define the functional contribution of NV cargo to anti-fibrotic remodeling, we compared Gene Ontology (GO) biological process and molecular function, together with KEGG pathway enrichment analyses, across the total NV proteome and proteins downregulated in activated fibroblasts following treatment with either direct NVs or silk-released NVs (**Table S16**). Comparative analysis identified 23 shared enriched terms across all three datasets, including 14 associated with actin cytoskeleton and actomyosin remodeling, such as actin filament organization, actin filament bundle assembly, actin cytoskeleton organization, and actomyosin structure organization (**Supplementary fig. S5D**). To further characterize proteins within these shared actin-associated pathways, we performed STRING protein–protein interaction network analysis on NV cargo proteins annotated to the GO term negative regulation of supramolecular fibre organization. This revealed a distinct cluster including CLASP1/2, PHLDB2, CAPZB, SPTAN1, SPTBN1, and ADD1/2/3, associated with cytoskeletal remodeling and actin filament regulation **(Supplementary fig. S5E)**. In addition, NV cargo contained LEMD3, a negative regulator of TGFβ signaling [73] (**Table S1**). Together, these data support the contribution of NV cargo proteins to actin cytoskeleton remodeling and fibroblast reprogramming.

### 3.6. In vivo application of silk-NV biomaterial in murine ischemia reperfusion injury model

To assess the therapeutic efficacy and biocompatibility of the silk-NV patch in an ischemia–reperfusion injury (IRI) model, we performed temporary ligation of the Left Anterior Descending coronary artery (LAD) to induce hypoxia, followed by reperfusion after 1 h by release of the loop. At the time of reperfusion, mice were left untreated (IRI) or received either a silk patch alone (IRI + empty patch) or an NV-loaded silk patch (IRI + silkNV) applied to the epicardial surface (**Fig. 6A**). Sham animals underwent the same surgical procedure, including heart puncture, without LAD ligation. B-mode echocardiography was performed at baseline, 24 hours post-IRI to confirm injury, and 28 days post-IRI to assess functional recovery (**Fig. 6A**). At 28 days, hearts were harvested and the left ventricles (LV) were dissected into three regions: the central infarct region for histological assessment of biocompatibility and fibrosis, and the LV apex and base for region-specific proteomic remodeling (**Fig. 6B**). Macroscopic examination of explanted hearts revealed no visible signs of inflammation, necrosis, or abnormal adhesions in either silk-treated group, supporting the overall biocompatibility of the patch (**Supplementary Fig. S6A**).

**Figure 6.**
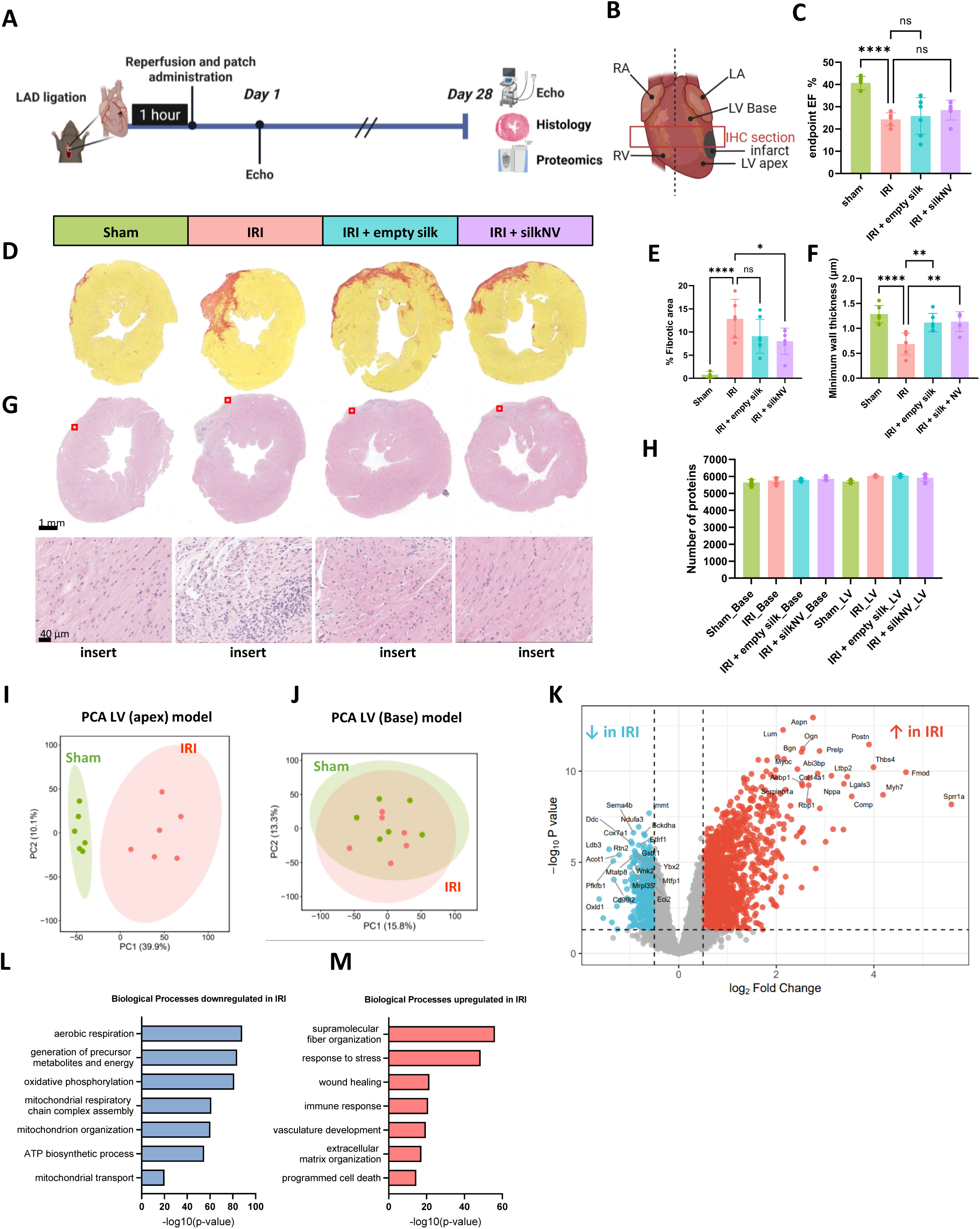
In vivo application of silk-NV biomaterial in murine myocardial infarction model. (**A**) Schematic of experimental workflow for Ischemia Reperfusion Injury (IRI) model and treatment with silk hydrogel patch alone of containing 5 µg of NVs. **(B)** Diagram of mouse heart dissection for analysis showing immunohistochemistry (IHC) section as well as left ventricule (LV) Base and LV apex for proteomics. **(C)** Bar plots of left ventricular ejection fraction (EF%) in treatment groups after 4 weeks of treatment. Sham n=6, IRI n=6, IRI+empty silk n=7 and IRI + silkNV n=6 (Dunnett’s multiple comparisons test, ***p < 0.0001, ns: not significant). (**D**) Picrosirius red staining of heart sections, with fibrotic tissue appearing pink and viable myocardium in yellow. (**E**) Quantification of infarct size expressed as percentage of fibrotic area relative to total myocardial area (Dunnett’s multiple comparisons test, *p < 0.05, ***p < 0.0001, ns: not significant). (**F**) Minimum wall thickness expressed in µm (Dunnett’s multiple comparisons test, **p < 0.01, ****p < 0.0001). (**G**) Hematoxylin-eosin (H&E) staining of heart sections. Red square indicates zoom insert. (**H**) Number of proteins identified in Base and LV samples for each treatment group. (**I-J**) Principal Component Analysis (PCA) of sham and IRI mouse left ventricular proteomes for apex region (**I**) and Base region (**J**). (**K**) Volcano plot comparing IRI vs sham control (t-test, adjusted p□<□0.05; Fold change >0.5 red — upregulated in IRI, Fold Change <-0.5 blue — downregulated in IRI. (K-L) Gene Ontology (GO) of biological processes (BP) identified in downregulated proteins (**L**) and upregulated proteins (**M**).

Left ventricular ejection fraction (EF) was quantified to evaluate cardiac function across treatment groups (**Fig. 6C**). At 28 days, all IRI groups exhibited significantly reduced EF compared to sham (24.3 ± 2.6% in IRI, 25.8 ± 7.6% in IRI + empty patch, and 28.5 ± 4.1% in IRI + silkNV vs. 40.7 ± 2.7% in sham), consistent with chronic ventricular remodeling and contractile dysfunction (**Fig. 6C**). Although silkNV treatment did not significantly improve EF, a trend towards increased EF was observed relative to the IRI group, which was more pronounced than in the empty silk group (**Fig 6C**). Importantly, electrocardiogram monitoring during surgery at the time of patch application showed no disruption to heart rate (data not shown), and echocardiographic assessment at day 28 revealed no differences in heart rate across treatment groups (**Supplementary Fig. S6B**). Furthermore, no adverse effects were observed in other cardiac functional parameters, including end-diastolic and end-systolic volumes, stroke volume, fractional shortening, cardiac output, and global longitudinal strain (**Supplementary Fig. S6B**), supporting that patch application did not further impair cardiac function.

To assess pathological remodeling and fibrosis, we performed Sirius Red staining on heart cross-sections and quantified the percentage of scar area (**Fig. 6D-E**). Fibrotic scar size was significantly increased in the IRI group compared with sham while treatment with NV-loaded silk significantly reduced fibrotic scar size relative to IRI (8.0 ± 2.6% vs. 12.8 ± 3.8%). A reduction was also observed in the empty patch group; however, this did not reach statistical significance (9.1 ± 3.4% vs. 12.8 ± 3.8%). Minimum LV wall thickness was quantified using the same sections, revealing a marked reduction in the IRI group compared with sham, indicative of substantial myocardial thinning, while both empty and NV-loaded silk significantly increased wall thickness relative to IRI (**Fig. 6F**). H&E staining showed reduced immune cell infiltration in both silk-treated groups compared to IRI alone at 28 days post-surgery, supporting the biocompatibility of the biomaterial and reduced inflammatory remodeling (**Fig. S6G**).

To further characterize global proteomic remodeling following IRI, quantitative proteomic analysis was performed across the four treatment groups and in two distinct regions: the LV apex (referred to as LV) and LV base (referred to as Base) (**Fig. 6B**). After filtering proteins with ≥70% valid values, an average of ∼5800 proteins were quantified per group, with consistent protein identification across all treatment conditions (**Fig. 6H**). Following imputation, a total of 6284 proteins were retained for downstream analysis (**Table S17**). Prior to treatment-group comparisons, model validation was performed by assessing sham versus IRI groups. To evaluate regional remodeling, principal component analysis (PCA) was conducted on LV apex and LV base samples from sham and IRI hearts separately. Distinct clustering of sham and IRI samples was observed in the LV apex (**Fig. 6I**), whereas partial overlap was evident in the LV base (**Fig. 6J**), indicating that proteomic alterations were more pronounced in the apical region. This spatial pattern is consistent with the LAD ligation site, with the apex located distal to the occlusion and the base proximal to it. We identified marked changes in protein expression between IRI and Sham groups at the LV-apex region, with 406 downregulated (FDR<0.05 and FC<-0.5) and 1324 upregulated proteins (FDR<0.05 and FC>0.5) (**Fig. 6K**). Gene ontology analysis revealed that downregulated proteins were mainly involved in metabolism regulation and mitochondria activity including oxidative phosphorylation and mitochondria respiratory chain complex assembly (**Fig. 6L, Table S18**). Upregulated proteins on the other hand were related to biological processes characteristic of a fibrotic remodeling including extracellular organization, immune response and programmed cell death (**Fig. 6M, Table S19**). To assess potential cardiotoxicity associated with silk patch treatment, we compared our dataset against a published panel of 12 doxorubicin-induced cardiotoxicity markers in mice[74]. Nine of these markers were detected in our dataset (ADIPOQ, SERPINA3N, RAB14, YWHAB, DYNLL2, KNG1, ACOT7, CAV1, MYH7), all of which remained unchanged across sham, IRI, and silk-treated groups, supporting the cardiac safety of the biomaterial platform (**Table S17**). Together these results validate our ischemia reperfusion injury model and support the biocompatibility of the silk-based patch in vivo.

### 3.7. NVs delivered via a silk hydrogel patch reprogram the left ventricular proteome toward repair following myocardial infarction

Understanding and isolating the effect of iPSC-NVs independently of the silk scaffold is essential for defining NV-specific molecular mechanisms of cardiac repair. Accordingly, multiple pairwise comparisons were performed (IRI vs sham, Empty_silk vs IRI, SilkNV vs IRI, and Empty_silk vs SilkNV). Proteins meeting the significance threshold (FDR < 0.1) are displayed in the heatmap in Figure 7A. Unsupervised hierarchical clustering revealed two major protein clusters that segregated predominantly by treatment group rather than surgical condition (**Fig. 7A**). Cluster 1 comprised proteins that were downregulated following IRI but selectively upregulated in the SilkNV-treated hearts, with expression levels closely resembling those observed in sham samples. Functional network analysis of cluster 1 revealed 4 main functional networks related to oxidative phosphorylation (i.e., NDUFB 2/3/4/6, COX6 and ATPK5), mitochondrial translation (i.e., MRPS 23/25/28/33/36 and TANGO2), sarcoplasmic reticulum (i.e., MLIP, TRDN, ALPK5 and CKM) and striated muscle contraction (i.e., TNNI3, SMIM20 and TPM1) (**Fig 7B**). Conversely, Cluster 2 contained proteins upregulated after IRI that were selectively downregulated toward baseline in the SilkNV group. Functional network analysis of cluster 2 revealed 3 main functional networks related to cell adhesion (i.e., FBLN1, APP and MAP1A), ECM remodeling (TIMP2, SERPINF1, MMP2 and THBS2) as well as regulation of renal sodium excretion (ACE and NPPB) (**Fig 7C**).

**Figure 7.**
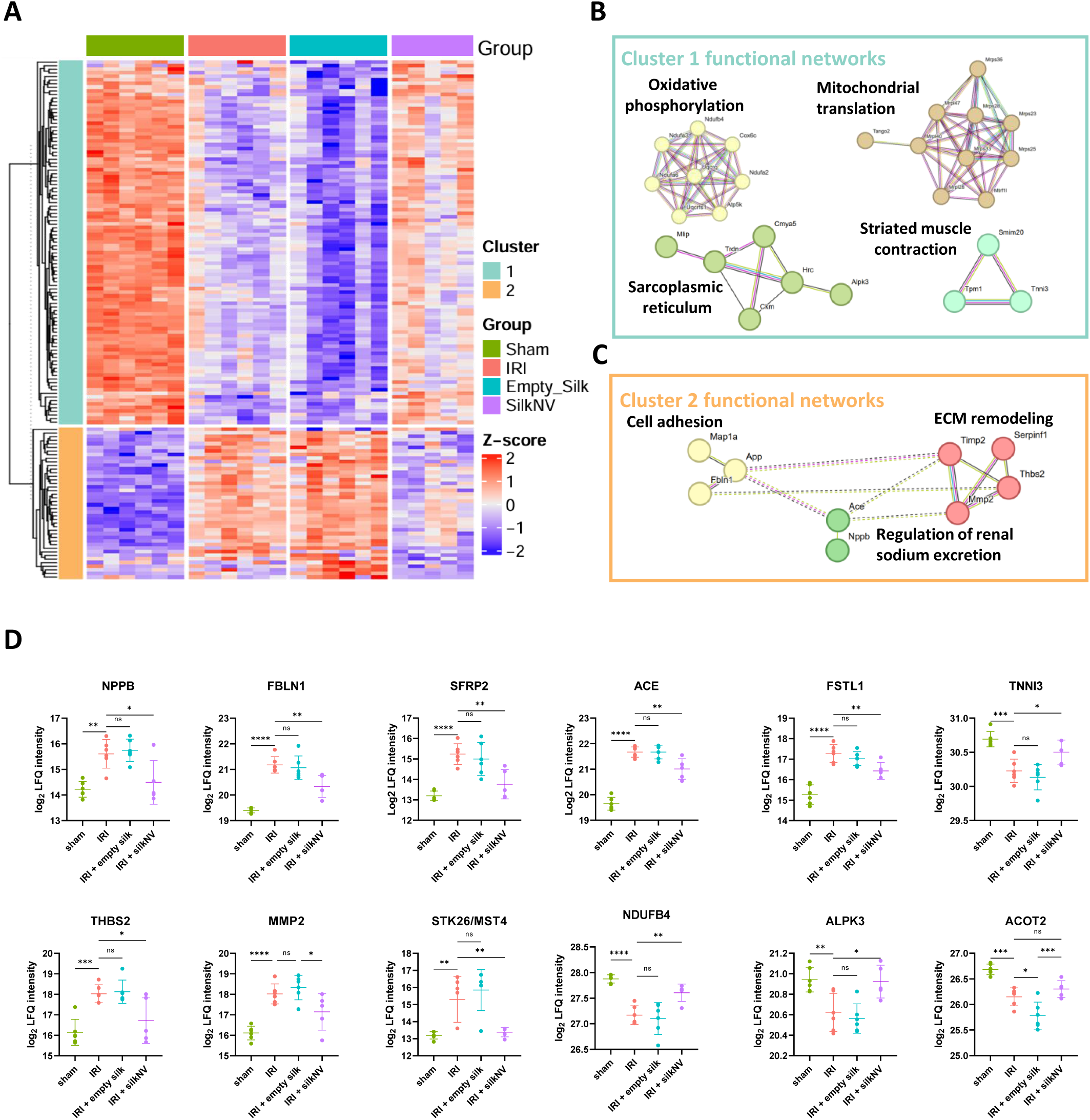
Nanovesicles delivered via a silk hydrogel patch reprogram the left ventricular proteome toward repair following myocardial infarction. **(A)** Heatmap of significantly regulated proteins across the four treatment groups (FDR < 0.1, limma multiple pairwise comparisons: IRI vs sham, Empty silk vs IRI, SilkNV vs IRI, and Empty silk vs SilkNV). (**B–C**) Functional network analysis of protein clusters identified from the heatmap, highlighting key protein interactions within cluster 1 (**B**) and cluster 2 (**C**) regulated by SilkNV treatment. (**D**) LFQ intensity of 12 clinically relevant heart failure–associated proteins significantly regulated by SilkNV treatment in vivo.

Notably, while the Empty_silk group resulted in a modest reduction in fibrotic area at the histological level, the proteomic profile largely mirrored or exacerbated IRI associated alterations across both clusters, suggesting limited proteomic modulation by silk alone. In contrast, SilkNV treatment was associated with a coordinated proteomic shift toward a sham-like profile across both clusters. We highlight 12 key recovery associated proteins that were specifically regulated by NV loaded silk but not by empty silk (**Fig. 7D**). The clinical heart failure biomarker NPPB was significantly upregulated in both IRI and IRI + empty silk groups compared with sham, whereas NV-loaded silk treatment reduced NPPB expression to near sham levels, consistent with reduced heart failure-associated molecular signatures. ACE, a therapeutic target of ACE inhibitors, a gold standard heart failure pharmacotherapy, was also specifically downregulated by NV treatment, consistent with cardioprotective remodeling. Proteins associated with cardiomyocyte function and structural integrity, including ALPK3 and TNNI3, were upregulated following NV-loaded silk treatment. Conversely, fibrosis and remodelling related proteins (MMP2, THBS2, FBLN1, SFRP2, and FSTL1) were significantly downregulated, consistent with attenuated pathological remodelling. Finally, metabolic recovery proteins involved in energy metabolism, including NDUFB4 and ACOT2, hallmarks of mitochondrial dysfunction in chronic heart failure, were also selectively modulated by NV delivery.

Collectively, these results demonstrate remodelling of the cardiac proteome 28 days following NV-loaded silk patch administration, supporting sustained NV-associated proteomic remodeling following IRI.

## 4. DISCUSSION

In this study we establish for the first time a di-tyrosine photo-crosslinked silk hydrogel platform for the encapsulation and sustained delivery of functional cell-derived NVs. Using iPSC-derived NVs generated by mechanical extrusion as a scalable EV-mimetic strategy, we demonstrate their efficient encapsulation into photo-crosslinked silk fibroin hydrogel, in turn, altering mechanical properties of the biomaterial, and retaining biological activity upon release both *in vitro* and *in vivo*.

Through extensive biophysical and proteomic characterization, we show that iPSC-derived NVs can be reproducibly generated at high yield and maintain a consistent bioactive cargo profile across preparations. When incorporated into silk hydrogels fabricated at microliter scale, NVs remain stable, retain their biophysical properties following release over 7 days, and remain detectable within the matrix for up to 28 days, supporting both sustained release and long-term local retention. Released NVs induce phenotypic, functional and proteomic changes in target cells both *in vitro* and *in vivo*, supporting the capacity of this biomaterial platform to provide localized and sustained bioactive delivery. We further demonstrate the tissue adhesive properties of the di-tyrosine silk hydrogel, enabling injection-free administration directly onto the heart surface in a myocardial infarction model, reducing scaring and promoting NV-driven remodeling of the cardiac tissue.

This system integrates three key advances: (i) the use of scalable iPSC-derived nanovesicles as an EV-mimetic platform, addressing limitations in EV manufacturing; (ii) a mechanically robust silk fibroin hydrogel formed via visible light-induced di-tyrosine crosslinking without chemical modification; and (iii) an injection-free, adhesive patch-based delivery strategy enabling localized and sustained therapeutic administration to the heart.

Cell-derived NVs have emerged as a compelling alternative to EVs, with comparable therapeutic potency, while addressing limited scalability of high-yield manufacturing of EVs with consistent therapeutic properties [18, 21, 54, 75–78]. A key advantage of mechanical cell extrusion is the rapid generation of nano-sized NVs in high yield. However, cell debris and cytosolic proteins generated during the mechanical extrusion could be potentially exposed on NV membranes and trigger immune response [79]. To limit the immunogenic risk, we employed a purification step to remove cytosolic/soluble factors generated during NV preparation. Previous studies within our research group have shown that NVs retain molecular cargo dependent on their cell source [80], can be modified through bioengineering [81], and can functionally regulate tissue repair processes, including promoting angiogenesis and cell survival, as well as reduction of fibrosis[54]. A standing question remains the reproducibility of extrusion-based NV generation from iPSC source. Here, we assess both technical consistency of NV generation (*intra-day* variation) and biological variation from cell preparations (*inter-day* variation). Biophysical characterization combined with proteomic profiling of NVs demonstrate high reproducibility in particle yield, size distribution, and proteome. Importantly, our data demonstrates that iPSC-derived NVs, enriched in their membrane composition relative to donor cells, retain important pluripotency markers and proteins implicated in regulation of tissue regeneration across and within preparations. These findings support the consistent, high-yield, and functional properties of NVs following their generation across different preparations [80, 82, 83] and from iPSC source [54].

Choosing an appropriate cell source is a critical consideration in the development of EV-mimetic therapeutics, as the biological origin of vesicles directly influences their molecular cargo, potency, and functional activity [84–88]. iPSCs offer several advantages including scalability through continuous expansion, potential for autologous sourcing, and broad therapeutic potential through anti-apoptotic, anti-inflammatory, pro-angiogenic, antioxidant, and anti-fibrotic properties [89–92]. Despite these advantages, vesicles from iPSC source are largely underrepresented in cardiac repair pre-clinical studies, with 64% of preclinical biomaterial-EV therapies for cardiac repair using MSCs-derived EVs, followed by endothelial progenitor- and endothelial-derived EVs being the next most commonly investigated [93]. This overrepresentation likely reflects the established regenerative profile and accessibility of MSCs but also highlights the limited exploration of alternative cell sources. A direct head-to-head comparison of different EV sources for cardiac repair determined that among the 6 cell line tested, ESCLEVs provided the best outcome after MI by reducing fibrosis and increasing angiogenesis [88]. In this context, induced pluripotent stem cells (iPSCs) may offer an attractive alternative, as they share key pluripotent and regenerative properties with ESCs while overcoming ethical limitations and offering patient-specific sourcing [94].

Systemic EV administration is limited by rapid immune-mediated clearance, poor tissue-specific targeting and the need for higher doses to achieve therapeutic concentrations at the site of injury [3, 95]. While NVs can be bioengineered to enhance tissue targeting [96–99], delivery route remains a critical determinant of efficacy and safety. Although using autologous IPSCs may reduce immunogenicity, systemic administration may still compromise their therapeutic utility by increasing off-target effects and dose requirements. Importantly, higher EV doses have been associated with cytotoxic effects, including impaired exocytosis and lysosomal activity [100], and increased MMP9 activity [101]. Therefore, controlling spatiotemporal EV delivery is crucial for optimal efficacy while avoiding potential side effects in non-target organs.

To address the challenge of systemic NV delivery, we incorporated NVs into a silk fibroin hydrogel system to control their spatiotemporal delivery directly to heart tissue. Silk fibroin, a natural biopolymer, is gaining traction for cardiovascular applications due to its biocompatibility, tunable porosity and degradation, low-cost, mechanical strength and intrinsic bioactivity, making it well suited to provide structural support to the infarcted myocardium compared to other natural biomaterials [45, 47, 102, 103]. Previous studies have demonstrated the feasibility of combining NVs in decellularized ECM-based [104] and hybrid polyvinyl alcohol/chitosan-based systems (PBA-CP) [105], enabling controlled release of NVs from different hydrogels, while maintaining wound healing capacity *in vivo*. Unlike ECM-based hydrogels, which often suffer from batch variability and limited tunability, or synthetic hydrogels which are largely biologically inert, silk combines reproducibility, bioactivity, and mechanical strength [45, 47]. Importantly, while previous studies have demonstrated the feasibility of incorporating vesicles into various natural and synthetic biomaterials for cardiac repair, these approaches remain limited by reliance on native EVs with limited scalability and invasiveness of intra-myocardial injection delivery [93]. In contrast, our study is the first to combine scalable iPSC-derived NVs with visible light photo-crosslinked silk fibroin hydrogels to achieve localized, sustained, and injection-free cardiac delivery.

Interestingly, NV incorporation altered the secondary structure of silk fibroin, increasing β-sheet content and reducing random coil structures, consistent with the observed increase in mechanical strength. This aligns with findings by Han et al., who reported that EV encapsulation in a silk fibroin hydrogel similarly increased β-sheet content and accelerated gelation time from 60 min to 5-10 min, supporting the concept of direct vesicle–silk interactions [51]. While the mechanisms remain to be determined, this may reflect interactions between tyrosine-containing NV surface proteins [106] and tyrosine-rich silk domains during di-tyrosine photo-crosslinking, or alternatively, NV surface properties acting as nucleation sites for β-sheet formation or promoting local silk chain rearrangement [45, 52, 53, 107, 108]. These matrix–vesicle interactions may also contribute to NV stabilization, as silk is known to preserve the structure and bioactivity of encapsulated biomacromolecules through conformational stabilization, hydrophobic shielding, and reduced aggregation at material interfaces[109]. Applied to NVs, such interactions may help maintain membrane integrity, preserve surface protein presentation, and protect bioactive cargo during encapsulation and release, thereby supporting sustained NV bioactivity and therapeutic efficacy. This bidirectional interaction between NVs and the silk matrix represents an important design consideration and warrants further investigation as a strategy to tune material properties, vesicle stability, and release kinetics.

The use of proteomic analysis is a major strength of this study, providing comprehensive molecular profiling to evaluate NV composition, recipient cell and tissue remodeling, and biomaterial safety both *in vitro* and *in vivo*. By integrating Gene Ontology analysis of the total NV proteome with the proteomic remodelling of activated cardiac fibroblasts following NV treatment, we identify convergent biological processes suggesting that NV cargo may attenuate fibroblast activation by limiting stress fiber formation and contractile remodeling associated with myofibroblast differentiation. Beyond mechanism, proteomics also enabled an unbiased safety assessment of the silk biomaterial, where comparison against established cardiotoxicity marker panels showed minimal perturbation of cardiotoxicity-associated proteins both *in vitro* and *in vivo*, supporting the biocompatibility of the silk-based delivery system [74, 110]. Importantly, *in vivo* proteomic profiling revealed that NV-loaded silk patches induced early molecular remodeling of the injured heart toward a reparative, sham-like state, restoring mitochondrial, contractile, and metabolic proteins while normalizing clinically relevant heart failure markers such as NPPB [111] and ACE [112]. These molecular changes were accompanied by reduced scar formation and preserved ventricular wall thickness, despite only modest ejection fraction recovery by echocardiography at 28 days. Similar temporal disconnects between molecular remodeling and functional recovery have been reported in other preclinical myocardial infarction studies, where proteomic changes were observed at earlier timepoints and preceded significant improvements in ejection fraction, which only became evident at 6 and 8 weeks post-treatment compared to 2 and 4 weeks [113]. Together, these findings highlight the unique value of proteomics not only as a discovery tool, but as a sensitive and translational framework to assess mechanism, safety, and therapeutic efficacy in the development of next-generation vesicle-based biomaterial therapies.

Several limitations of this study should be acknowledged. Our *in vivo* assessment was performed exclusively in male mice and given the known sex-specific differences in cardiac remodeling and therapeutic response after myocardial infarction, future studies should evaluate the efficacy of this platform in both sexes. In addition, further optimization of NV release kinetics through better understanding of NV-silk interaction mechanisms, therapeutic dosing, and timing of administration will be important to define the optimal therapeutic window and maximize regenerative outcomes.

## 5. CONCLUSION

In conclusion, we demonstrate that adhesive silk fibroin hydrogels provide an effective platform for the localized and sustained delivery of therapeutic nanovesicles to the heart. By combining the scalability and bioactivity of stem cell-derived NVs with the mechanical strength, adhesive, and biological properties of silk, this platform enables both structural support and prolonged molecular modulation of cardiac repair processes. We show that sustained NV release preserves anti-fibrotic bioactivity, modulating integrin-mediated signalling, actomyosin organisation, and cell–matrix adhesion in cardiac fibroblasts, while promoting molecular remodeling and reducing the fibrotic scar of the injured myocardium *in vivo*. Importantly, our findings reveal a previously underappreciated bidirectional interaction between NVs and the silk matrix, where vesicle incorporation influences biomaterial structure and mechanics, highlighting new opportunities to optimize vesicle retention, stability, and release. Together, this adhesive silk-NV platform represents a novel acellular therapeutic strategy with strong translational potential for spatiotemporally controlled, tissue-targeted delivery in cardiac repair and broader regenerative medicine applications.

## Supplementary Figure Legends

**Supplementary Figure S1. Generation and proteomic profiling of iPSC-derived nanovesicles**

(**A**) Representative particle size distribution of iPSC-NV determined by nanoparticle tracking analysis (NTA). (**B**) Schematic overview of SP3 (single-pot, solid-phase-enhanced) sample preparation for LC-MS/MS analysis and data processing for iPSCs, iPSC-derived NVs and HUVECs (non-stem control). (**C**) Gene Ontology (GO) of cellular compartment (CC) identified in top 100 most abundant proteins in iPSC-NV proteome. (**D**) Venn diagram of surface annotated proteins in iPSC-NV proteome, using in silico surfaceome SURFY [66], cell surface protein atlas CSPA [67], and cell surface protein annotation (GO:0009986) from UniProt [68]. Of the surface-annotated NV proteins identified (by at least 2/3 databases), 57/183 proteins contained tyrosine (Y) residues in their peptide sequences. (**E**) Principal Component analysis of iPSC (cell), iPSC (NV) and HUVEC (cell) proteome.

**Supplementary Figure S2. Function and remodeling of TGF-treated cardiac fibroblasts by iPSC-derived NVs**

(**A**) Representative immunofluorescence microscopy images of human Ventricular Cardiac Fibroblasts (hVCFs) treated with either PBS (vehicle control), TGFβ (10 ng/mL), or TGFβ+NV (30µg/mL) for 72 hours and stained with anti-αSMA antibody and Hoechst (nuclei),scale bar = 500μm. (**B**) αSMA positive area (*n* = 3, mean ± SEM. * *p* < 0.01). (**C**) Volcano plot comparing TGFβ vs PBS (control) treated hVCFs (t-test p□<□0.05; red – upregulated, blue – downregulated in TGFβ). (**D-E**) Selected Gene Ontology (GO) terms enriched in (**D**) upregulated proteins in TGFβ vs PBS (and those unique to TGFβ) (**E**) and downregulated proteins in TGFβ vs PBS (and those unique to PBS); CC: Cellular Component, BP: Biological Process, MF: Molecular Function.

**Supplementary Figure S3. Additional characterization of silk hydrogels**

**(A)** Degradation profile of silk hydrogels without (2% silk) and with NVs (2% silk +NV) expressed as percentage of mass loss (%) over time (*n* = 5, mean ± SEM. 2-way ANOVA* *p* < 0.05, *** *p*<0.0005). **(B)** Representative images of silk hydrogels without (2% silk) and with NVs (2% silk +NV) with corresponding autofluorescence associated with di-tyrosine bonds. (C) Diagram of injection-free administration of 5 μL of silk hydrogel followed by 3 min of light exposure (n=3). (**D**) After crosslinking, hearts were excised and ran through flowing water to assess patch adherence to pericardium.

**Supplementary Figure S4. Sustained release of intact and functional iPSC-derived NVs from silk fibroin hydrogels**

(**A-G**) Size distribution of released NVs and empty hydrogel control as determined by Nanoparticle Tracking Analysis (NTA) from day 1-7. Dotted line indicates empty silk control and full line is for releasates from silk-NV composite.

**Supplementary Figure S5. Silk-NV composite reprogrammed cardiac fibroblast proteome**

(**A**) Heatmap of significantly downregulated proteins (one-way ANOVA) across TGFβ (activated), TGFβ+2% silk (empty silk), and TGFβ+2% silk_NV (NV-loaded silk). Highlighted cluster includes proteins specifically downregulated in the NV-loaded group. (**B**) Gene Ontology (GO) enrichment of proteins in the NV-specific downregulated cluster; CC: Cellular Component, BP: Biological Process, MF: Molecular Function, REAC: Reactome. (**C**) Selected GO terms enriched in proteins significantly upregulated in TGFβ+2% silk_NV vs 2% silk (and uniquely present in TGFβ+2% silk_NV); KEGG (Kyoto Encyclopedia of Genes and Genomes) pathways. (**D**) Venn Diagram of intersection of GO terms between NV proteome and downregulated proteins in NV and silkNV-treated fibroblasts. (**E**) STRING network of NV proteome-associated proteins annotated to the GO term negative regulation of supramolecular fibre organization.

**Supplementary Figure S6. Biocompatibility of silk-NV biomaterial in murine myocardial infarction model**

(**A**) Macroscopic (gross) examination of the left ventricle after 28 days to assess inflammation, necrosis, or abnormal adhesions. (**B**) 28-day echocardiogram analysis of key cardiac functional parameters, including end-diastolic (EDV) and end-systolic volumes, stroke volume, fractional shortening, cardiac output, and global longitudinal strain (GLS%).

