## Supplementary Data (Figures S1-6) for "Adhesive silk hydrogel patches for localized and sustained delivery of cell-derived nanovesicles"

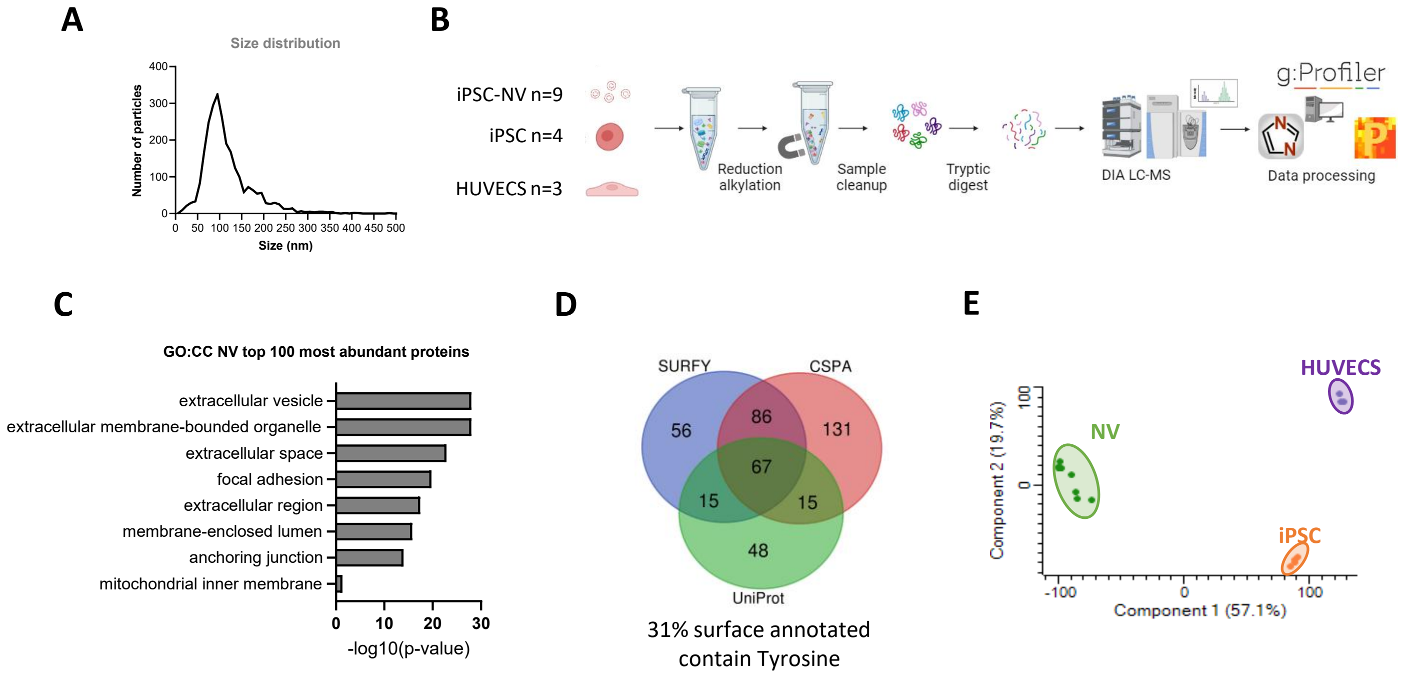


**Supplementary Figure S1. Generation and proteomic profiling of iPSC-derived NVs**

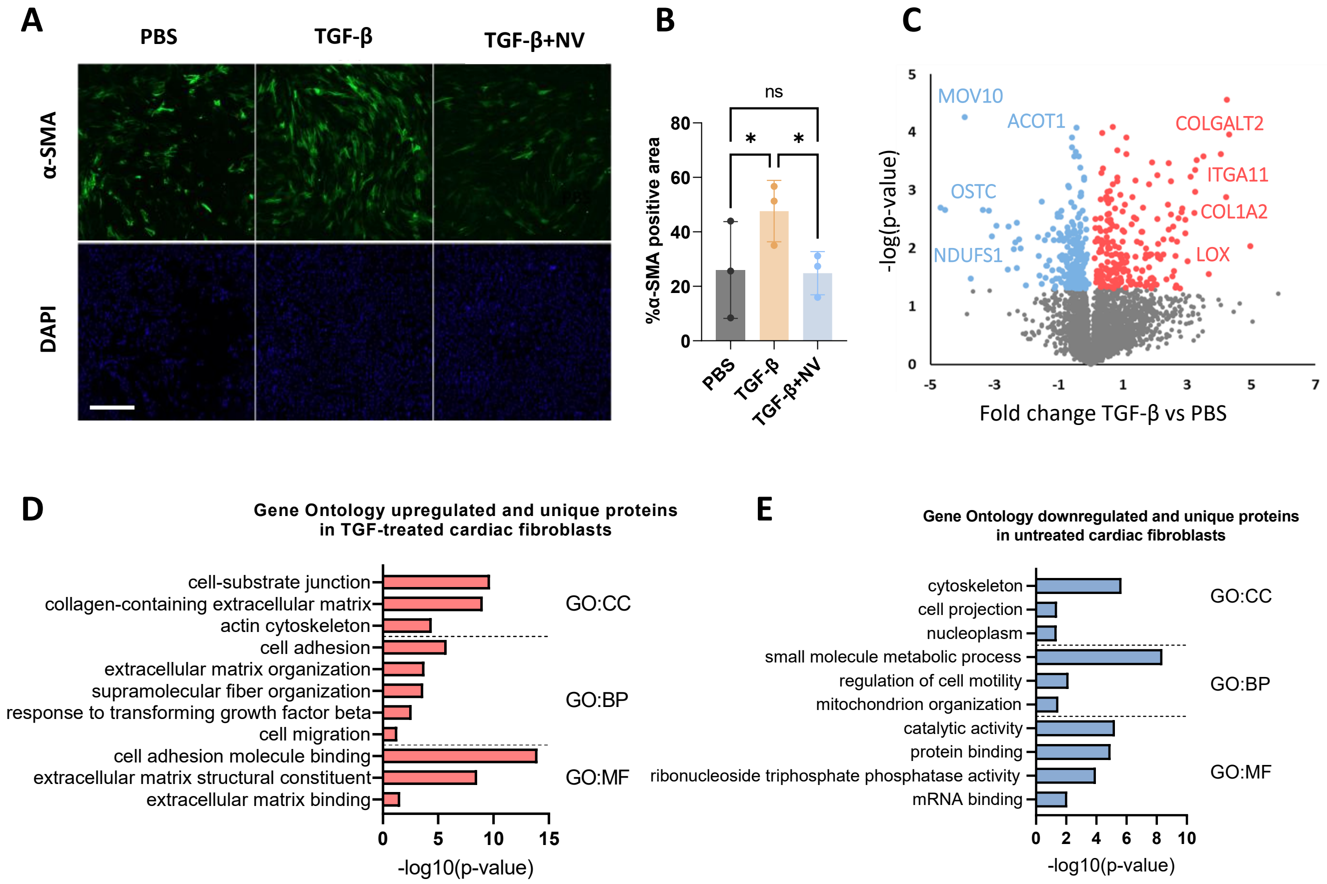


**Supplementary Figure S2. iPSC-derived NVs reverse TGFβ-driven cardiac fibroblast remodelling**

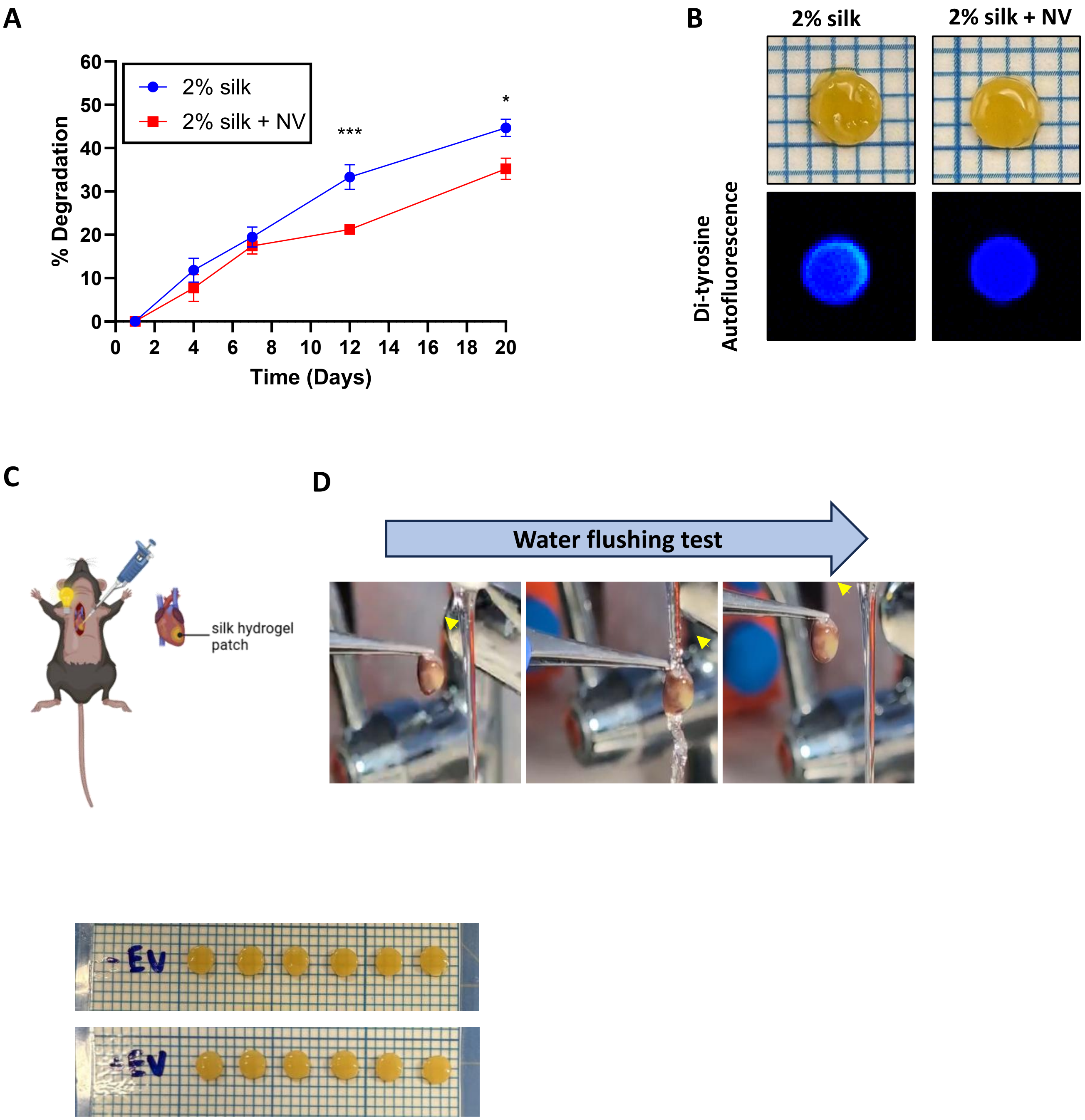


**Supplementary Figure S3. Additional characterization of silk hydrogels**

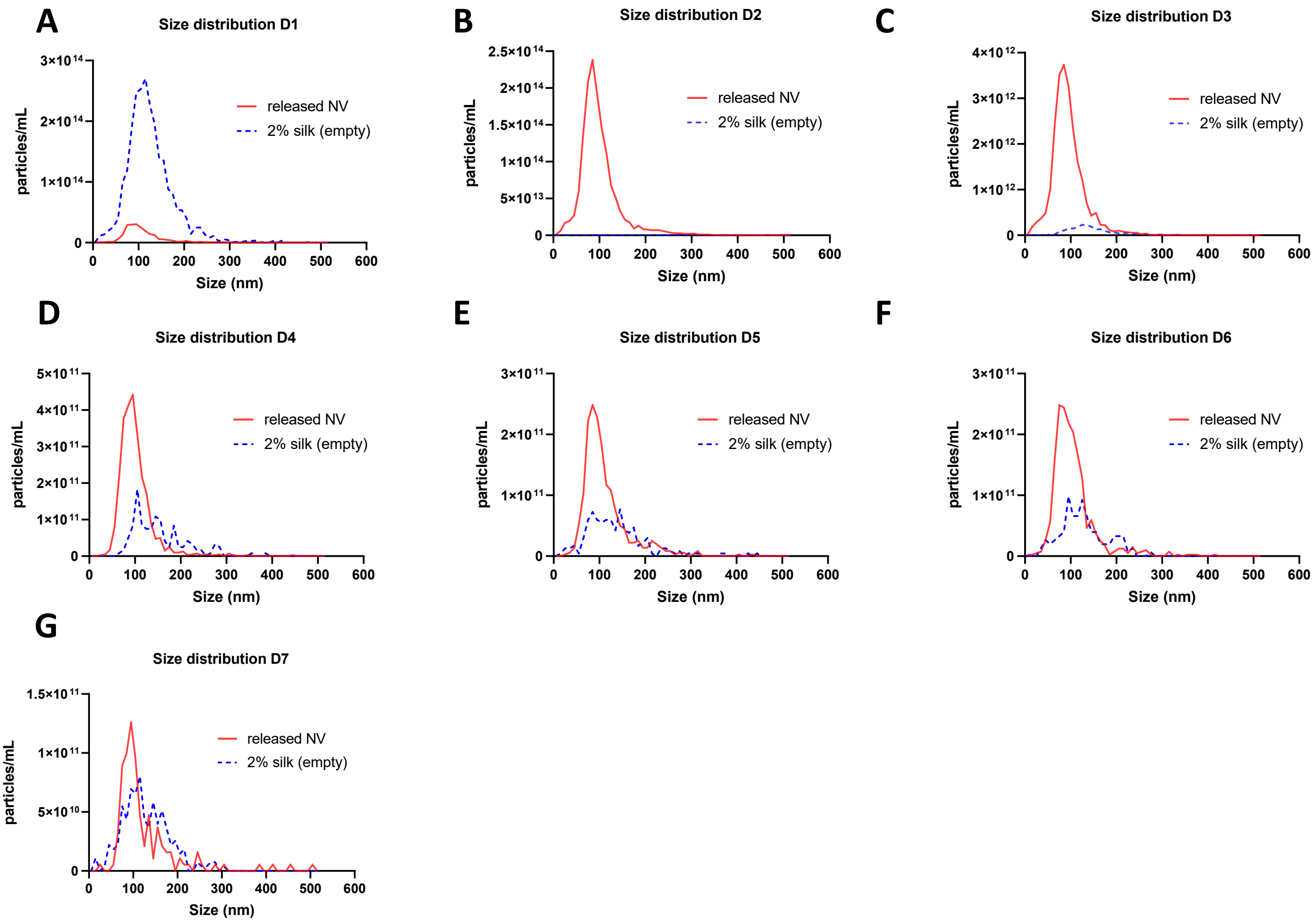


**Supplementary Figure S4. Sustained release of intact and functional iPSC-derived NVs from silk fibroin hydrogels**

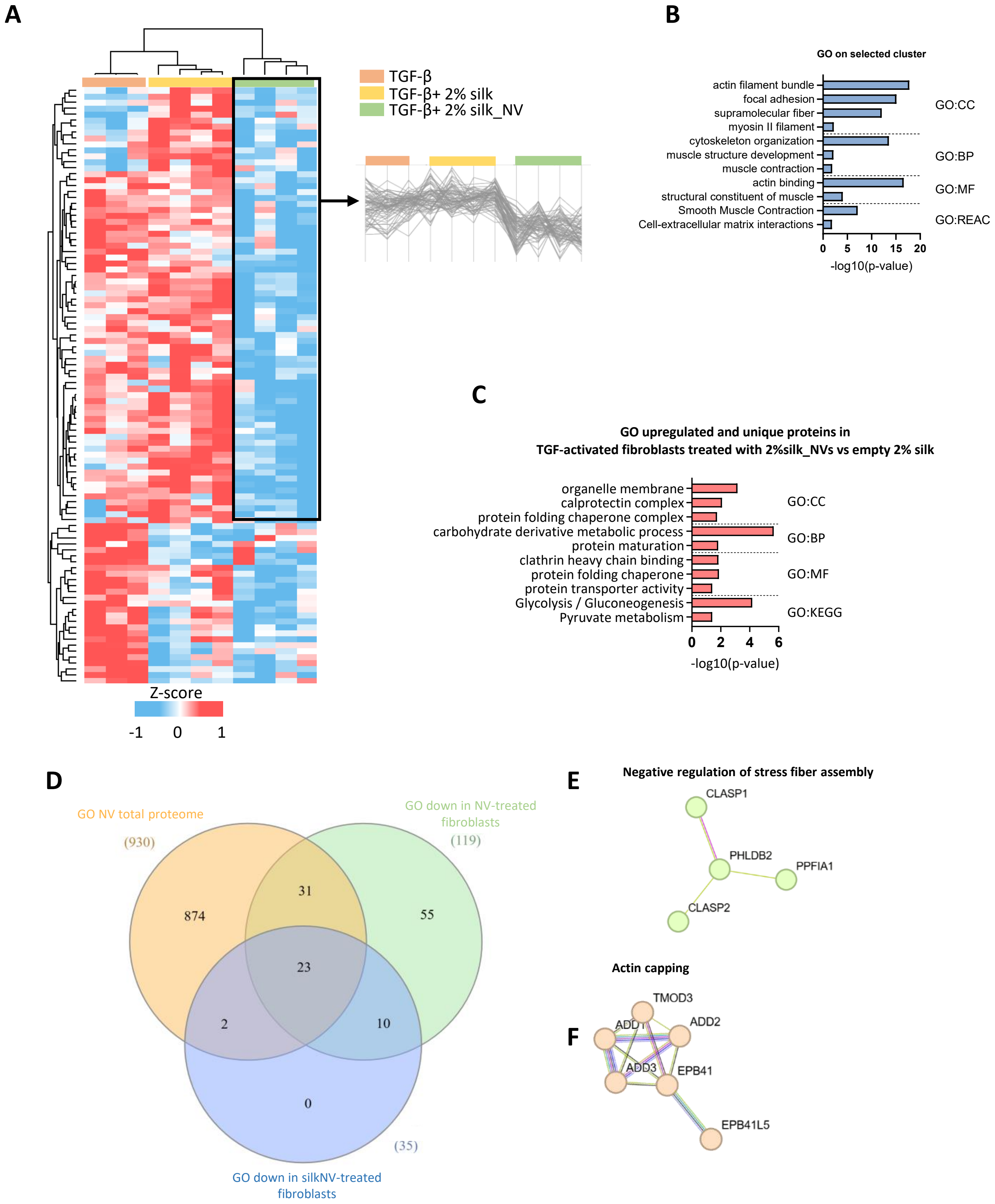


**Supplementary Figure S5. Silk-NV composite reprogrammed cardiac fibroblast proteome**

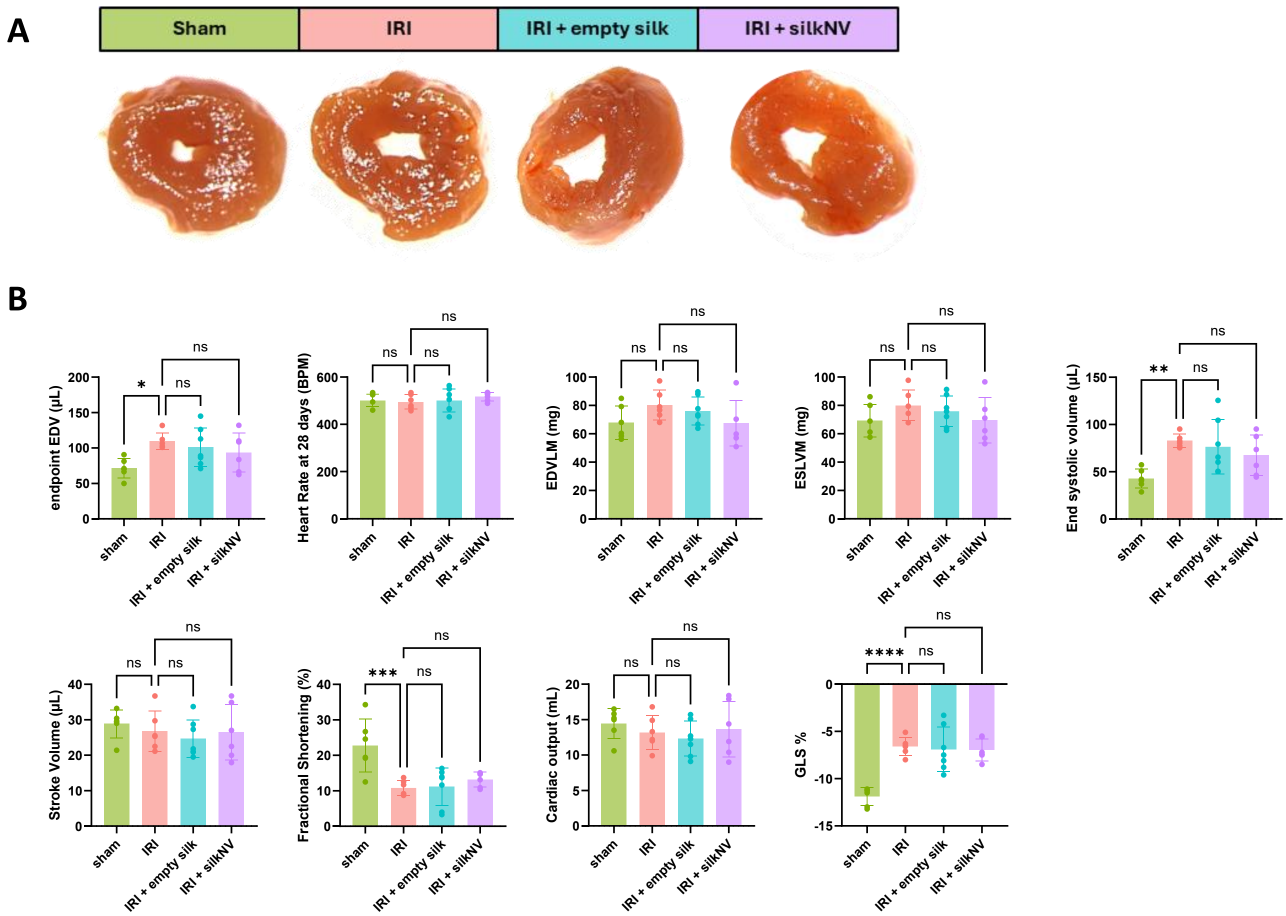


**Supplementary Figure S6. Biocompatibility of silk-NV biomaterial in murine myocardial infarction model**
